# Reading specific memories from human neurons before and after sleep

**DOI:** 10.1101/2025.07.01.662486

**Authors:** Yuanyi Ding, Soraya L. S. Dunn, John J. Sakon, Zahra M. Aghajan, Christopher A. Chan, Aditya Datta, Chenda Duan, Yipeng Zhang, Joel I. Berger, Ariane E. Rhone, Kirill V. Nourski, Hiroto Kawasaki, Matthew A. Howard, Vwani P. Roychowdhury, Itzhak Fried

## Abstract

The ability to retrieve a single episode encountered just once is a hallmark of human intelligence and episodic memory[1]. Yet, decoding a specific memory from neuronal activity in the human brain remains a formidable challenge. Here, we develop a transformer neural network model[2, 3] trained on neuronal spikes from intracranial microelectrodes recorded during a single viewing of an audiovisual episode. Combining spikes throughout the brain via cross-channel attention[4], capable of discovering neural patterns spread across brain regions and timescales, individual participant models predict vocalization of specific concepts such as persons or places during out-of-distribution memory recall. Brain regions differentially contribute to memory decoding before and after sleep. Models trained using only medial temporal lobe (MTL) spikes significantly decode concepts before but not after sleep, while models trained using only frontal cortex (FC) spikes decode concepts after but not before sleep, with better FC decoding relating to increased non-REM sleep. These findings suggest a system-wide distribution of information across neural populations that transforms over wake/sleep cycles[5]. Such decoding of internally generated memories suggests a path towards brain-computer interfaces to treat episodic memory disorders through enhancement or muting of specific memories.

---

Reading internal cognitive states of the human mind from neural activity, such as specific episodic memories, remains a major challenge for contemporary neuroscience. Emerging approaches involve training deep learning neural network models on electrophysiological activity recorded while participants engage with external stimuli and subsequently uncovering the stimuli from held-out neural data, such as decoding objects in an image[6, 7], speech[8, 9], heard music[10, 11], or characters in a video[12–14]. Methods that decode internal cognitive states[15], however, require many presentations of static images [16, 17] or words[18] to form a training set and typically pool neural data across several participants[13, 18]. A neural mechanism to uncover internal thoughts could use concept cells, neurons that selectively respond to concepts such as persons or places regardless of their sensory input[19], and internally reactivate prior to free recall[20]. Indeed, selective responses from concept neurons can reliably predict memory retrieval of a concept[20, 21] and decode visual inputs using linear methods[16]. Given the sparse coding by MTL concept neurons[22], however, the ability to identify and potentially target neurons selective to a given concept remains limited—particularly for new experiences[13, 23]. Optogenetic methods that selectively target cells, such as engram cells[24], remain unavailable in humans. Further, models using single neuron inputs typically require laborious manual curation after data collection[25], restricting the ability to decode from previously-trained models in real-time.

In sum, decoding of specific memories from one-shot experience based on neural responses remains infeasible with current methods. Yet, the brain can successfully assimilate a single experience into durable memory traces extending from the hippocampal-entorhinal system to the neocortex[26]. This distributive process occurs prominently during non-rapid eye movement (NREM) sleep, where offline consolidation of hippocampus-dependent declarative memories occurs through coordinated hippocampal replay and hippocampal–neocortical oscillations[5, 27]. We hypothesize, then, that the distribution of episodic information throughout the brain can support a decoding model of specific memories during awake recall, and that these representations and the resultant decoding will change following a night of sleep.

To determine if we can identify retrieval of concepts encoded during a single episodic experience, we asked 15 participants undergoing clinical monitoring with intracranial electrodes for intractable epilepsy[28] to watch a single 42-minute episode (from the TV series “24”) and subsequently: 1) perform five minutes verbal free recall of episode content from memory, 2) distinguish 75 “old” video clips (0.5-3.0 s) of the watched episode from 75 “new” clips of an unseen episode in the same season of the “24” TV series, and 3) verbally answer six questions, each accompanied by a still image for 45 seconds each (Fig. 1a). We term the last memory test “cued free recall”, as the question and still image provide a cue but the participants free recall related content. As we anticipate neural representations to change across initial encoding, subsequent recall, and recall after sleep[5], we repeated the memory tests following overnight sleep. The tests indicate participants encoded the episode well (Fig. 1d), correctly answering 77.3 ± 8.5% of old/new recognition memory questions (Fig. 1d, right; d-prime=1.65 ± 0.61).

**Fig. 1.**
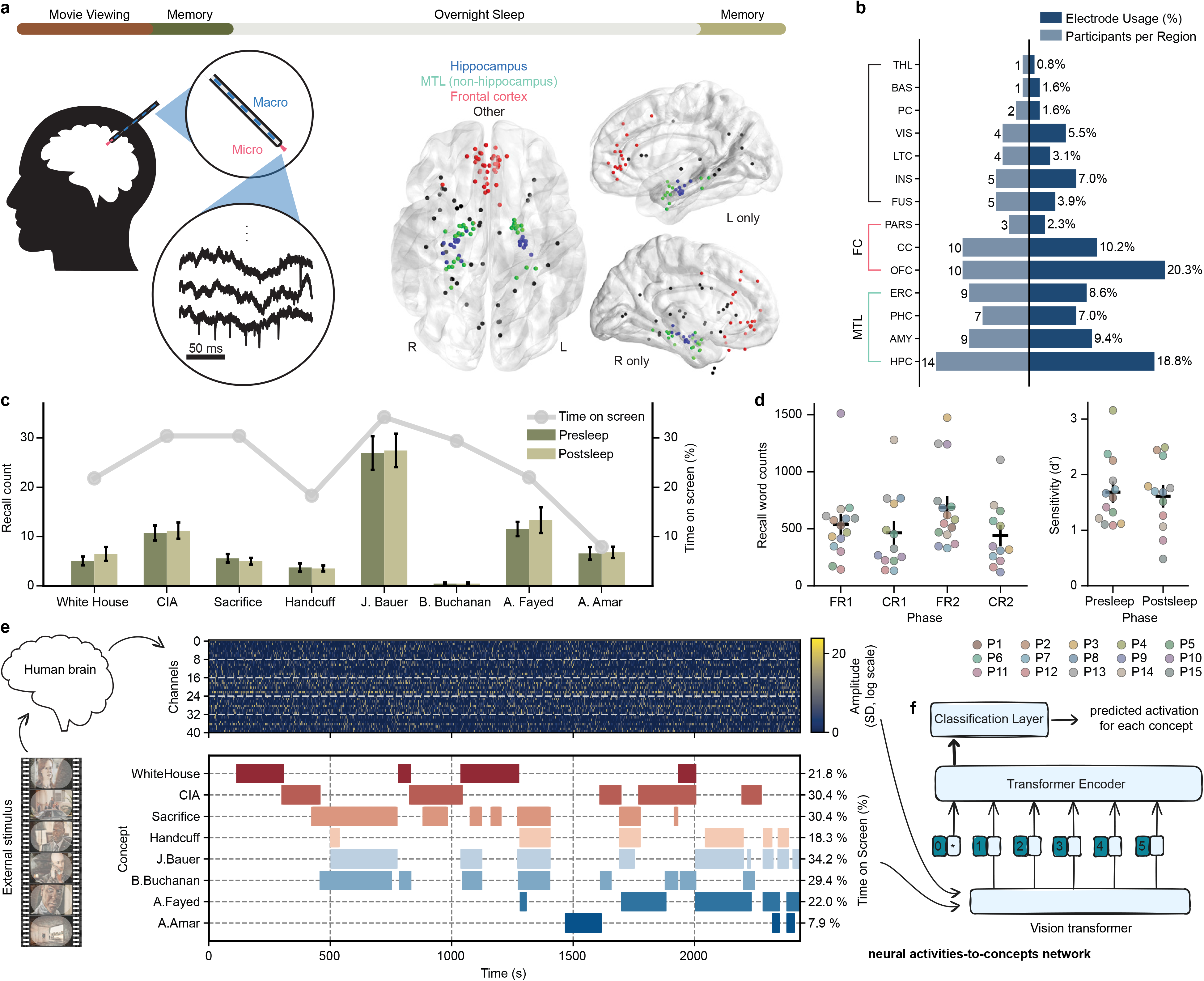
Methods and behavioral results. **a**, Top: 15 participants watched an episode of the show 24 followed by several memory tests. After sleep, they completed the same set of memory tests the next morning. Left: Throughout the entire experiment, we recorded brain activity using Behnke-Fried depth electrodes with microwires protruding from the macroelectrode into medial brain structures. Center and right: Localization of microwire recording sites across all participants superimposed on a standard brain. Center: ventral view, anterior towards top. Right: same microwire sites superimposed on mid-sagittal views of left and right hemisphere. **b**, Microwire distribution across brain regions. Bars extending to the right indicate the proportion of electrodes in each region, while bars extending to the left indicate the number of unique participants with electrodes in each region. Abbreviations: HPC (hippocampus), AMY (amygdala), PHC (parahippocampal cortex), ERC (entorhinal cortex), OFC (orbitofrontal cortex), CC (cingulate cortex), FUS (fusiform gyrus), INS (insular cortex), LTC (lateral temporal cortex), PARS (pars orbitalis), VIS (visual cortex). **c**, Average number of vocalized recalls for each concept collected during the presleep and postsleep recall tasks (bars) contrasted with the %time the concept was shown onscreen in the episode (lines). Error bars (black) reflect standard error (SE) across 15 participants. **d**, Total words spoken (left) during free recall (FR) and cued free recall (CFR) before and after sleep (1 vs. 2) shows that participants reliably recall far more than target concepts. Recognition memory sensitivity (d-prime, right, *N* = 12 participants), which assesses true positive vs. false positive accuracy of old/new clips, confirms reliable memory for movie content before and after sleep. **e**, (Top): example raster plot of positive-polarity spike amplitudes across the movie episode. Gray dashed lines separate bundles of eight microwires. The input matrix has shape 2 *× channels × bins*, with the first dimension indexing spike polarity (see Methods). (Bottom): Occurrence of concepts during episode. **f**, Schematic of the encoding model. We trained a vision transformer to learn the mapping between neural activity and the presence of semantic concepts shown in the external stimulus (TV episode). The model aims to identify consistent neural representations associated with each concept across different moments in the movie.

As 15 participants performed the tasks before and after a night of sleep, we recorded stable neuronal activity (Extended Data Fig. 1) from 73.1 ± 19.6 microwires per patient—capable of recording single unit activity[28, 29]. Microwires typically localized to medial brain regions—particularly Medial Temporal Lobe (MTL) and Frontal Cortex (FC)(Fig. 1a-b). We thresholded signals from all microwire channels to extract clusterless voltage spike trains[30] for each electrode (Methods). This method allows rapid, real-time decoding of new data[31, 32], making our models potentially more compatible with future BCI devices by foregoing spike sorting. We designed a transformer neural network model[2] that computes channel-wise self-attention within each bundle of eight microwires and then combines information across bundles via cross-channel attention[4] (Extended Data Fig. 2a; see Methods). Training of models occurs separately for each participant, with model stopping determined in four early participants (Extended Data Fig. 3), using the clusterless voltages recorded while participants watched a single presentation of the episode. We hypothesize the model can then identify neuronal patterns related to sets of concepts, including persons (e.g., the main character J. Bauer), places (e.g., the White House), and abstract concepts (e.g. sacrifice) present in the episode (Fig 1e). We chose eight concepts to test based on the amount of time onscreen (Fig. 1d-e) and the tendency of participants to recall them (Fig. 1d).

## Memory decoding performance

To predict the identity of concepts in out-of-distribution data recorded during recall (pooled free and cued free recall vocalizations where available) we inference from each participant’s model trained on viewing period data. This process outputs a model activation between 0 and 1 for each concept independently over the course of each recall period (Fig. 2a), indicating how strongly the neural pattern at each moment during internally generated memory retrieval matches the signature for that concept acquired from model training. For concepts where models successfully identify associated neuronal patterns, we expect increases in model activation prior to vocalizations of that given concept, thereby reflecting memory reinstatement. To assess such reinstatement, we developed an evaluation metric termed Memory Confidence Score (MCS) (Fig. 2b; Methods), which quantifies the model activation prior to vocalization of a given concept vs. the activations for that concept before non-concept vocalizations (Fig. 2a, bottom). To calculate MCS, first we average the area above baseline -4 to 0 seconds before each vocalization of a given concept (*A*_real_, Fig. 2b, left), where the baseline represents mean activation across the whole recall period for that concept (Fig. 2a, dashed line). We then take an equal number of randomly-sampled vocalizations for other words and repeat this process 1500 times to create a surrogate set of mean concept activations over baseline ({*Asurrogate*^(*k*)^}*_k_*_=1_^1500^, Fig. 2b, middle). The MCS represents the percentile rank for the real value vs. all surrogates values (Fig. 2b, right; Fig. 2c, bottom). This way, we determine the MCS for a concept not from the absolute value of model activity, but rather by comparing the statistics of the model output for vocalizations of a specific concept against those of surrogate vocalizations. We limit the evaluation of MCS to concepts vocalized at least three times during recall (Fig. 2d).

**Fig. 2.**
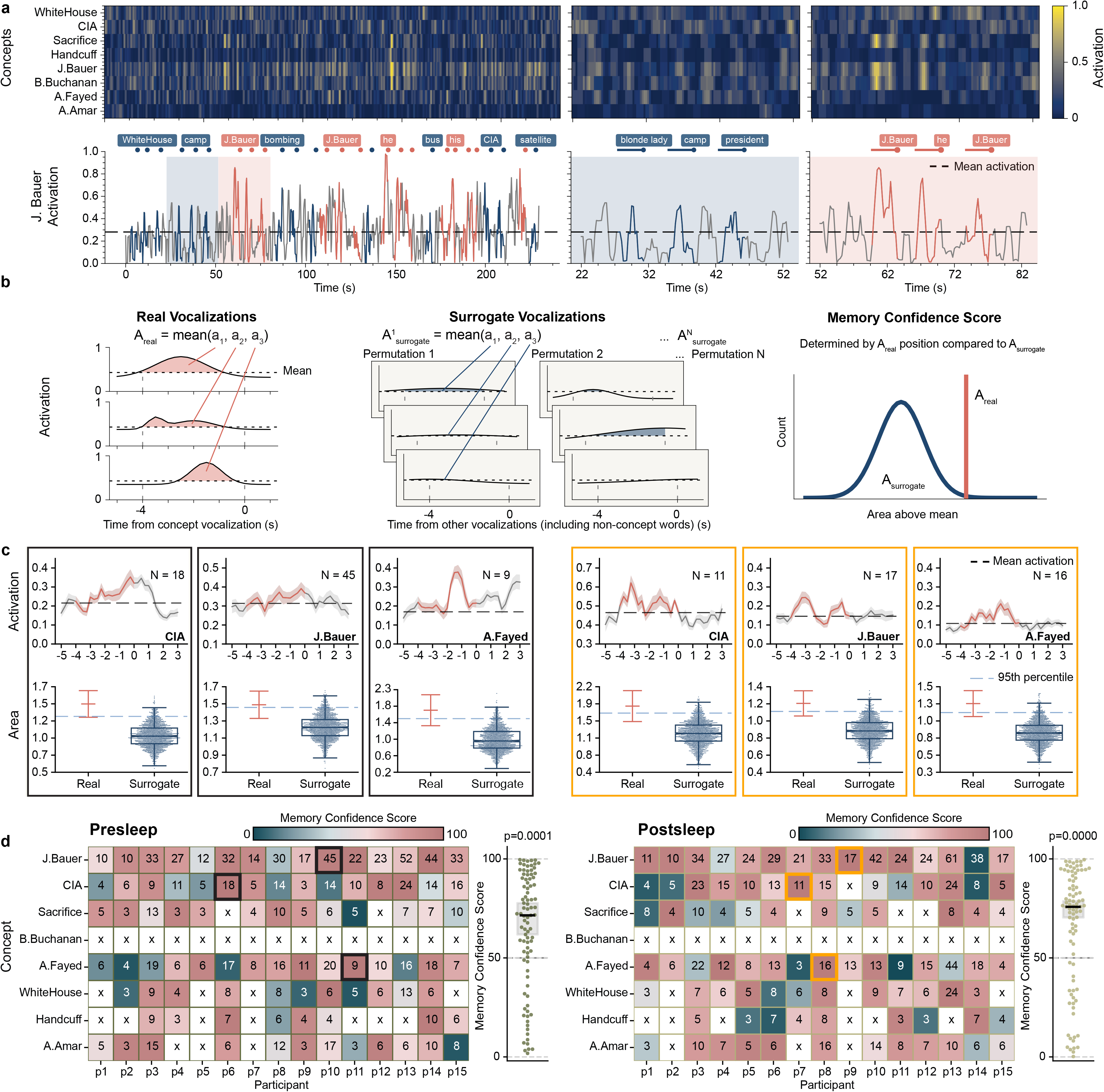
Memory Confidence Score (MCS) unveils decoding of memories prior to vocalization. **a**, An example model output from a single participant illustrates the MCS procedure. The top-left heatmap shows model activations for all concepts during a 230-second presleep free recall session. Below, a line plot traces the activation of the concept ‘J. Bauer’ over time. Red dots show vocalizations referring to the main character ‘J. Bauer’, blue dots show vocalizations to other concepts, and colored line segments represent 4-second intervals preceding each vocalization onset. The panels in the middle and right figures, zoomed in views of the blue and red shaded sections in the left figure, show stronger activation patterns prior to ‘J. Bauer’ vocalizations compared to unrelated vocalizations. **b**, To quantify the degree of activations prior to related concepts, we compute the area under the activation curve above the mean baseline within each 4-second pre-vocalization window. In the toy example, for a concept mentioned three times (left), the average of these areas defines *A*_real_. We then randomly sample three unrelated vocalizations, compute the same metric, and repeat this process *N* = 1500 times to generate a surrogate distribution *A_surrogate_* = {*A_surrogate_*^(1)^,…,*A_surrogate_*^(1500)^} (middle). We define the MCS as the percentile rank of *A*_real_ within this surrogate distribution. **c**, Six example concepts with high MCS using our Full model, three from presleep recall (left) and three from postsleep recall (right). The top plot of each pair displays the mean model activation (±SE) aligned to vocalization onset across all occurrences, with the red segment indicating the 4-second pre-vocalization window used for evaluation. The bottom plot visualizes the percentile ranking of *A*_real_ within the surrogate distribution *A*surrogate. Real denotes the mean and SE of the activation areas contributing to *A*_real_, while Surrogate shows the distribution *A*_surrogate_ as a swarm plot. **d**, Heatmaps of population-level MCS across all participants and concepts for presleep (left) and postsleep (right) recall for our Full model. Each colored cell in the heatmap represents a concept (row) mentioned *≥*3x by a participant (column); numbers indicate the total number of vocalizations for that concept by that participant. Concept decoding compared to chance (50%) shows significant effects for both conditions (presleep: *p* = 1.03 *×* 10*^−^*^4^, postsleep: *p* = 4.52 *×* 10*^−^*^5^; Wilcoxon signed-rank), indicating consistent decoding of concepts across participants. The three black outlined cells (left) represent the presleep examples in **c** and the three orange outlined cells (right) represent the postsleep examples in **c**.

Across all 15 participants, our model shows significant decoding of concepts during presleep recall with pooled MCS significantly above chance (*Z* = 3.88*, p* = 1.03 × 10*^−^*^4^*, D* = 0.458; Wilcoxon signed-rank test and one-sample Cohen’s D for all MCS vs. 50% comparisons; Fig. 2d). We show several examples of significantly decoded concepts for single participants in Fig. 2c. In addition, our model significantly decodes the population of concepts during postsleep memory tests (*Z* = 4.08*, p* = 4.52 × 10*^−^*^5^*, D* = 0.492), indicating robust memory decoding performance. Do we achieve optimal decoding just prior to vocalization, as expected if our model decodes memories? Running identical model inference and MCS metrics as in our main results but on stepwise windows before or after the -4 to 0 s window used in Fig. 2 shows decreasing decoding in both directions (Fig. 3c, left), suggesting that our method unveils internal generation of memories reinstated just prior to vocalization. We also achieve significant decoding for concepts when using models that separately inference on only free recall (Extended Data Fig. 4a) or cued free recall (Extended Data Fig. 4d), indicating our model can independently decode internal memories generated in both recall tasks, but pool them in upcoming analyses to increase sample size for tests with stricter demands.

**Fig. 3.**
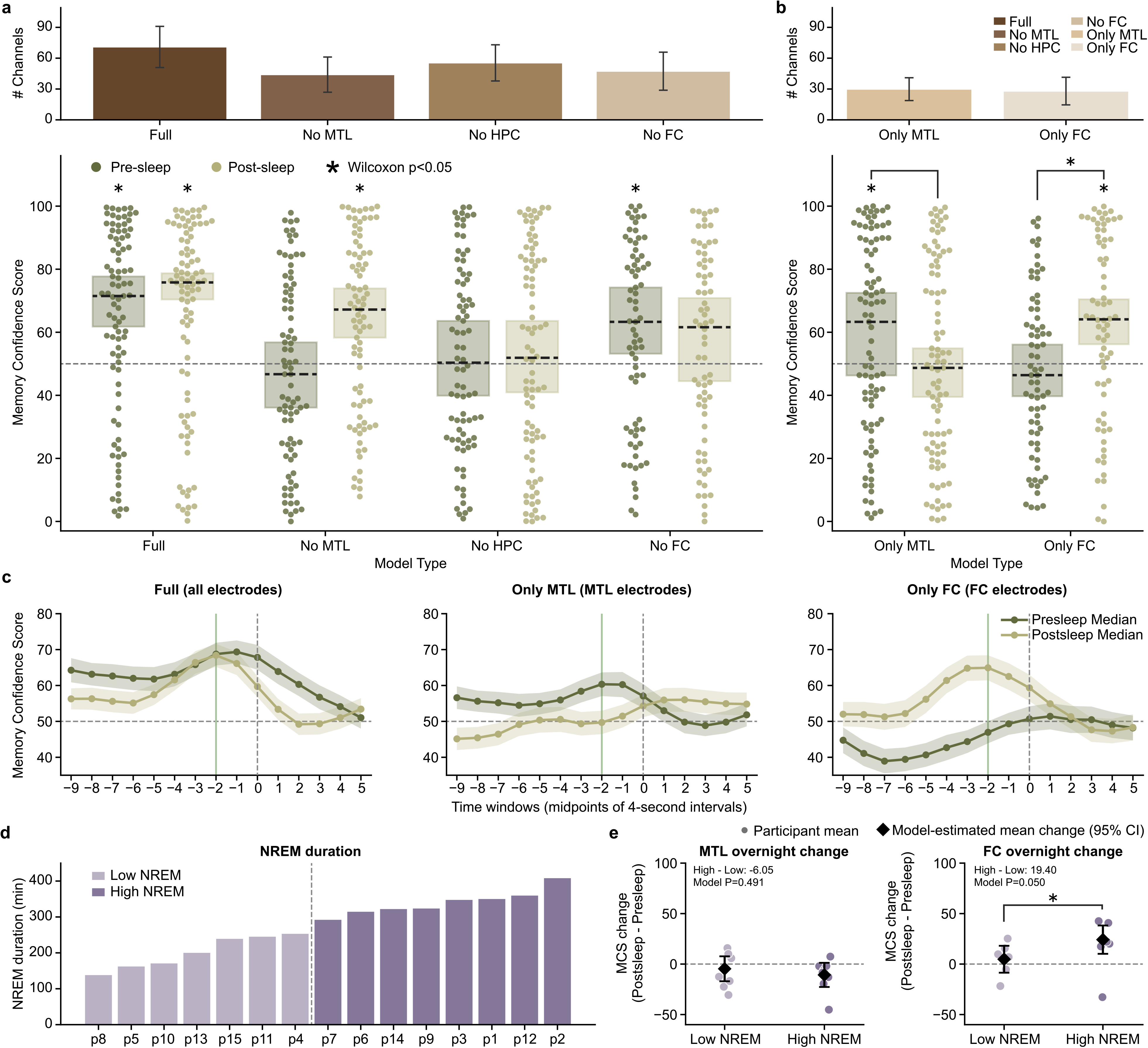
Medial temporal lobe (MTL) and frontal cortex (FC) differentially contribute to pre- and postsleep memory decoding. **a**, **Top**: mean*±*SE channel counts for each model; **Bottom**: Presleep and postsleep Memory Confidence Scores for the Full model (same data as Fig. 2d) vs. models with removal of clusterless voltage spike data recorded in MTL (including hippocampus), hippocampus (HPC), and FC. Each dot shows a single concept decoded for the given model trained on an individual participant’s data, dashed black lines show median, rectangles show interquartile range, and asterisks indicate significance of a Wilcoxon signed-rank test for concepts grouped across participants vs. chance (50% decoding). Note that the Full and MTL models include all (N=15) participants, while the No HPC (N=14) and No FC (N=12) models test only those participants that originally possess those regions (Extended Data Table 1). **b**, Same conventions as **a**, but for models restricted to channels localized in MTL (Only MTL, N=15) and FC (Only FC, N=12). **c**, Median pooled MCS across all participants, calculated from every four-second window centered from -9 to +5 seconds relative to concept vocalization onset with shaded areas indicating *±*1 s.e.m. For example, the -2 s point in these plots (green line) inferences on neural data in the window -4 to 0 s from vocalization (all MCS reported prior to this plot have used this -4 to 0 s window). The Full model (left), Only MTL model (center), and Only FC model (right) all show peak decoding in windows centered 1 to 2 seconds prior to vocalization for both pre- and postsleep. **d**, Using the duration of NREM sleep (left) for each participant, we compare overnight change in MCS for low vs. high NREM sleep groups for the Only MTL (center) and Only FC (right) models. Reported estimates and *p* values compare the high and low NREM groups using fixed effects from mixed effects models with participant-level random intercepts.

We run several control models to show the robustness of our deep learning decoding approach. First, to ensure our model does not reflect coincident overfitting, we run 100 surrogate models after circularly shifting microwire channel inputs by a random amount of time in the movie viewing period for each model (Methods). For both presleep and postsleep, all 100 surrogate models show lower median MCS than the correctly-aligned data (presleep: mean *Z* = −0.058, mean *D* = −0.006; postsleep: mean *Z* = 0.200, mean *D* = 0.021; Extended Data Fig. 5). Second, logistic regression decodes concepts significantly worse than our transformer-based model in both presleep and postsleep (presleep: *Z* = 2.970*, p* = 0.0030*, D* = −0.331, postsleep: *Z* = 3.840*, p* = 1.234 × 10*^−^*4*, D* = −0.440; paired Wilcoxon test and Cohen’s D for paired samples for all paired tests; Extended Data Fig. 6a), suggesting deep learning takes advantage of non-linear and spatiotemporal neural population interactions[13, 33]. Third, we attempt a similar transformer decoding framework using broadband oscillatory features in macrowire iEEG data[34], which covers more brain regions but at worse spatial resolution, instead of clusterless voltage spikes. The iEEG-trained model also decodes significantly worse than our clusterless-trained model for both presleep and postsleep (presleep: *Z* = 2.886*, p* = 0.0039*, D* = −0.337, postsleep: *Z* = 3.001*, p* = 0.0027*, D* = −0.331; Extended Data Fig. 6b). Fourth, to test sensitivity to surrogate pool composition (Fig. 2b), we recalculate MCS using vocalizations of either all other modeled concepts (Extended Data Fig. 7a) or, more stringently, only other concepts with MCS above chance level decoding from the Full model (Fig. 2d) within the same participant and recall period (Extended Data Fig. 7b). MCS remained significantly above chance for both controls during presleep (*p* = 0.0005 and *p* = 0.0016, respectively) and postsleep recall (*p* = 0.0002 and *p* = 0.0010; Wilcoxon signed-rank tests for each). Fifth, to test whether our approach can decode more than just recalled memories, we use the same participant-specific models trained on viewing to predict concept identity for participants with complete recognition memory test data (n = 12, Methods) as they viewed repeated video clips from the episode. Using model predictions within a two-second window centered on the midpoint of each annotated clip, we measure MCS using the same framework as Fig. 2d) by comparing clips containing a given concept with concept-absent clips. We find significant decoding of concepts across our participant population during presleep (*p* = 0.0053) and postsleep (*p* = 3.15 × 10*^−^*5; Wilcoxon signed-rank tests for each), suggesting our model can also identify neural patterns representing concept information during repeated clip viewing.

We also perform several analyses to characterize the effectiveness of our model. First, we determine if highly correlated concepts—in other words, those that share a high proportion of screen time in the video—show similar levels of decoding. Taking each pair of concepts for each participant, we relate the correlation of their continuous model predictions throughout each presleep or postsleep recall period (Extended Data Fig. 9b) to the absolute difference in their MCS. We find no relation between these metrics during presleep (*β* = −3.35±7.12, *p* = 0.638) or postsleep (*β* = 2.98±7.10, *p* = 0.675; linear mixed effects models; estimates are *β* ± SE), suggesting the decoding we show across our population (Fig. 2) does not rely on only a subset of related concepts. Second, we ask if concepts with more vocalizations show better MCS. Plotting the average MCS for each participant vs. the average number of recalls (a rough proxy for recall ability, Extended Data Fig. 10), we find little correlation between them presleep (*R* = 0.01*, p* = 0.98) and a trend towards correlation postsleep (*R* = 0.39*, p* = 0.16), with little correlation when relating the overnight change in MCS with recalls (*R* = 0.18*, p* = 0.53). Considering recall sample size has little relation with decoding of a given concept, our results do not appear driven by well-recalled concepts. Third, having used four participants to tune our model stopping hyperparameter (Methods), we measured the population MCS for the remaining 11 participants to confirm our model works for the remainder. Indeed, we find significant concept decoding across these other participants for both pre- and postsleep (*p* = 0.0053 and *p <* 0.0001; Wilcoxon signed-rank tests; Extended Data Fig. 11). Finally, while we developed our model to use voltage amplitudes from microwires without clustering for use in future real-time decoding applications (see Discussion), we ask if it performs better than a model using manually-sorted spikes. Comparing their MCS directly (Extended Data Fig. 12), the clusterless model shows stronger but not significantly different decoding presleep (*Z* = 0.820*, p* = 0.4120*, D* = 0.068) but does decode significantly better than sorted spikes postsleep (*Z* = 2.170*, p* = 0.0300*, D* = 0.214). These results suggest spikes removed by manual spike sorting (i.e., the noise cluster) or the voltage amplitudes themselves possibly contain information that contributes to our clusterless model.

## MTL and FC differentially contribute to memory decoding before and after sleep

To evaluate the contribution of specific brain regions on the performance of our memory decoder, we trained several additional models that excluded channels based on electrode localization (Fig. 1a, right and Extended Data Table 1). These models include removal of MTL (No MTL, which excludes microwires in hippocampus, amygdala, parahippocampal cortex, and entorhinal cortex), hippocampus (No HPC), and FC (No FC, which excludes microwires in orbitofrontal, superiorfrontal, inferiorfrontal, and anterior cingulate cortex). For each model we then measure the MCS both before and after sleep—using the same process shown in Figs. 2 and 3—to assess whether we can decode concepts without the contribution from the removed region.

For the 14 participants with microwires localized to both MTL and at least one other region (since a participant without MTL coverage must have remaining channels to run a model), the No MTL model fails to decode concepts above chance level in presleep memory tests (Fig. 3a; *Z* = −0.740*, p* = 0.4593*, D* = −0.077) but successfully decodes them in postsleep (Fig. 3a; *Z* = 3.691*, p* = 2.23 × 10*^−^*^4^*, D* = 0.445). For the 14 participants with microwires localized to both hippocampus and at least one other region, the No HPC model provides little evidence for concept decoding above chance level for memory tests both presleep (Fig. 3a; *Z* = 0.546*, p* = 0.5854*, D* = 0.056) and postsleep (Fig. 3a; *Z* = 0.579*, p* = 0.5625*, D* = 0.060). For the 12 participants with microwires localized to both FC and at least one other region, however, the No FC model successfully decodes concepts above chance level for memory tests presleep (Fig. 3a; *Z* = 2.452*, p* = 0.0142*, D* = 0.296) but only moderately postsleep (Fig. 3a; *Z* = 1.644*, p* = 0.1001*, D* = 0.196). These results suggest MTL—particularly hippocampus—provides crucial inputs to the full model both before and after sleep, while FC contributes primarily during postsleep.

Considering that removal of MTL and FC channels impact our full model in different ways, we ask if models trained using only MTL (Only MTL) or only FC (Only FC) can decode concepts during recall. Extended Data Table 1 summarizes the number of microwires from each region, with all 15 participants possessing at least 16 microwires in MTL and 12 participants possessing at least 16 microwires in FC. We find that the Only MTL model successfully decodes concepts above chance level for memory tests presleep (Fig. 3b; *Z* = 2.387*, p* = 0.0170*, D* = 0.254) but only weakly postsleep (Fig. 3b; *Z* = 0.104*, p* = 0.9169*, D* = 0.006), albeit with no significant difference when comparing the MCS distributions directly (*Z* = 1.327*, p* = 0.1846*, D* = 0.170). The Only FC model, meanwhile, fails to significantly decode concepts above chance level during presleep memory tests (Fig. 3b; *Z* = −0.742*, p* = 0.4581*, D* = −0.094) but does significantly decode them during postsleep (Fig. 3b; *Z* = 3.029*, p* = 0.0025*, D* = 0.392), with a significant difference when comparing them directly (*Z* = 2.373*, p* = 0.0176*, D* = 0.332). As with the Full model, both the Only MTL and Only FC model decoding peaks just prior to vocalization, as anticipated for recall of concepts (Fig. 3c). Combined, the Only MTL and Only FC models represent a double dissociation between pre- and postsleep decoding (Region × Period interaction: z=3.2, p=0.0016, mixed effects model), with Only MTL capable of significantly decoding concepts before sleep and Only FC capable of decoding concepts after sleep, but not vice versa, suggesting consolidation of specific concept memories from MTL to FC over one night of sleep.

If consolidation during overnight sleep shifts concept representations from MTL to FC, we expect participants with more NREM sleep to show a greater MTL-to-FC shift [5, 27]. We measure the duration of NREM sleep for each of our 15 participants using sleep scoring from iEEG [35] (Extended Data Fig. 3d). Splitting participants into high vs. low NREM groups, we find a significant effect between pre- vs. postsleep decoding for the Only FC (mean change 19.4 MCS, *p* = 0.05) but not the Only MTL (mean change -6.05, *p* = 0.49) model (Fig. 3e), with a trend in the interaction between them (mean change 25.45, *p* = 0.054; Extended Data Fig. 13a-d). We also directly relate NREM sleep with the overnight change in MCS for each model using mixed effects models (Extended Data Fig. 13e). While Only MTL shows little change from pre- to postsleep decoding (-2.1 MCS/hr, slope parameter from mixed effects model, *p* = 0.53), we find a trend for high NREM sleep leading to stronger FC decoding (6.0 MCS/hr, *p* = 0.094) and stronger FC relative to MTL change (8.1 MCS/hr, *p* = 0.098). Further, we recorded from a 16th participant that performed the same protocol as the first 15 but started with the video watch and memory tests at noon and completed the second set of memory tests 9.5 hours later without sleeping (iEEG confirmed N=0 hours NREM). Adding this participant to our previous analyses (Extended Data Fig. 13f-g), we find significantly greater FC and FC relative to MTL decoding for the high NREM group (Only MTL, mean -7.74, *p* = 0.36; Only FC, mean=21.4, *p* = 0.025; FC vs. MTL, mean=29.1, *p* = 0.022) and a significant association between NREM sleep and MCS decoding with the mixed effects model for both FC and FC relative to MTL (Only MTL, -2.7 MCS/h, *p* = 0.28; Only FC, 5.6 MCS/h, *p* = 0.035; FC vs. MTL, 8.4 MCS/h, *p* = 0.022). Overall, our results support the notion that the shift from MTL to FC decoding overnight occurs due to NREM-dependent sleep consolidation.

## Decoding not related to neuron selectivity

Does the extraction of specific memories achieved by our model depend on the recruitment of cells specifically responsive to concepts comprising those memories? In the human MTL, concept cells frequently respond to characters in visual media[19, 20]. We therefore investigate to what degree neuronal firing from potential concept cells evoked by the presence of characters in the video contributes to our transformer decoding performance. We hypothesize that if character-selective units provide the primary source of information for our models then removing them from the model input should severely degrade the MCS for many concepts. Therefore, we generated frame-level labeling to precisely measure on-screen character presence for 10 main characters that map onto five of our decoded concepts (Methods). Taking 536 manually sorted single- and multi-units with a minimum firing rate of 0.5 spikes/second over the video watch period, pooled across all 15 participants (Fig. 4a, Extended Data Table 3), we use a permutation t-test to identify units that significantly increase in firing rate when a given character appears onscreen (Fig. 4b-d; Methods). We find 52 units from 14 participants that selectively respond (*p <* 0.01 permutation t-test) to a single character (Fig. 4e; Extended Data Fig. 14a). To determine if our models rely on this character tuning for decoding, for each of these 14 participants we evaluate the model with all microwire channels that possess character-selective neurons removed (3.6±2.7 mean±std channels/participant removed) and calculate the MCS for the same set of concepts as before (Fig. 2b). Relating the MCS for each concept within each participant for the channel-removal model with the corresponding MCS for the Full model, we find no significant change in decoding performance for the channel removal model across our population for both presleep and postsleep (Fig. 4f; presleep: mean MCS change = -2.58, *Z* = −0.785*, p* = 0.463*, D* = −0.279; postsleep: mean MCS change = -3.27, *Z* = −1.852*, p* = 0.068*, D* = −0.601). Performing the corresponding character-responsiveness analysis additionally including 8 neurons that respond significantly to multiple characters yields similar results (Extended Data Fig. 14d). To make a matched comparison, for each participant we also evaluated 60 controls where for each we remove a random subset of channels equal to the number of channels removed as before, while matching the distribution of removed channels within electrode bundles. E.g., since P1 has 4 character-selective neurons from 4/48 channels, each of the 60 controls for this participant removes a random 4/48 channels—excluding channels containing character-responsive neurons—while preserving the same channels-per-bundle pattern as the character-selective channel removal. Comparing the character-selective channel removal model with the matched random removal model, we find no evidence removing character-selective channels harms decoding more than equivalent random channels (Extended Data Fig. 15a vs. 15b; presleep: *β* = 0.39 ± 2.19, *p* = 0.859; post-sleep: *β* = −0.89 ± 1.70, *p* = 0.602; mixed effects models; estimates are *β* ± SE). In sum, these results suggest that character-responsive units do not centrally contribute to how our model decodes concepts during recall.

**Fig. 4.**
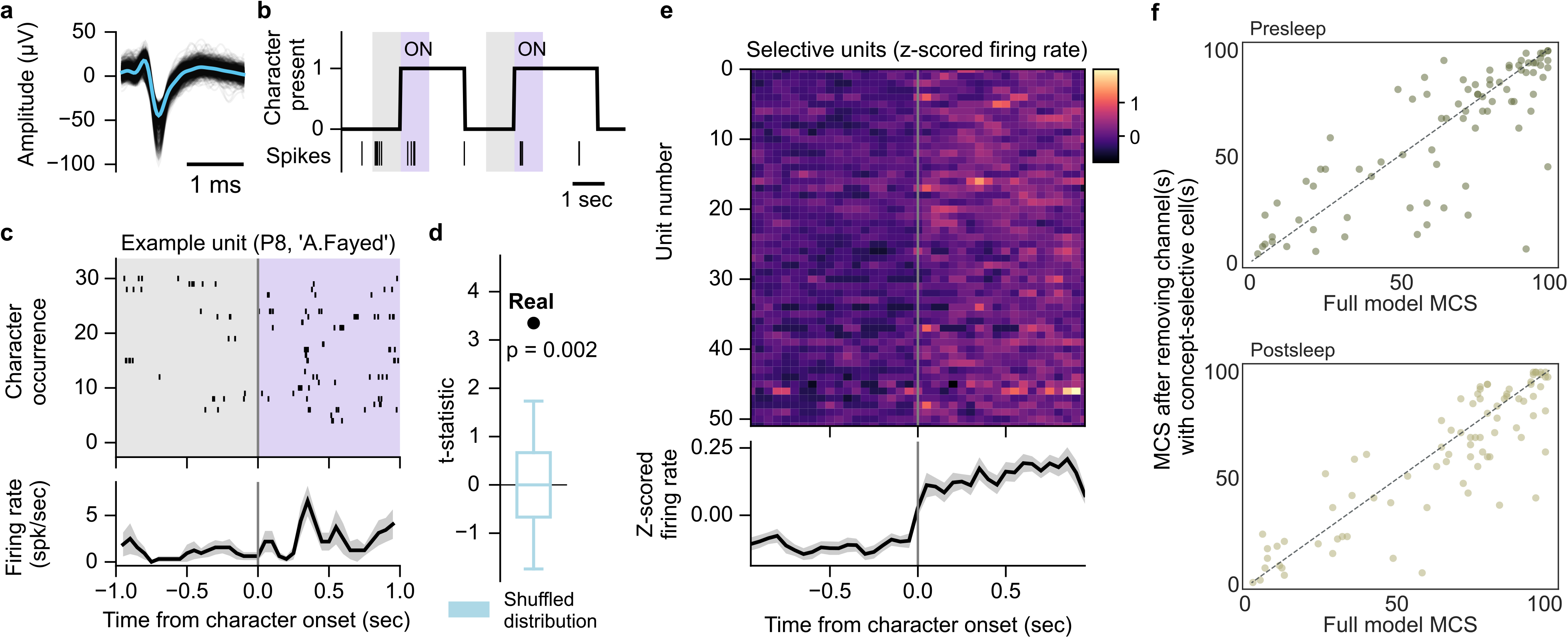
Character-selective cells cannot explain decoding performance. **a**, Waveforms of a manually spike sorted single unit (black lines) overlaid with the mean waveform (blue line). **b**, Example labeling for presence of character ‘A.Fayed’, with spiking activity of an example character-selective unit (unit shown in (**a**)) shown as vertical ticks below. Shaded regions indicate analysis windows 1 second prior to character onset (gray) and 1 second post-onset (purple). **c**, Spiking activity and peri-stimulus time histogram (PSTH) of the example unit in (**a**) over 51 appearances of ‘A.Fayed’. PSTH (bottom) black line shows mean firing rate across appearances in 100 ms bins with 50 ms overlap, gray shade represents SE. **d**, Observed t-statistic (‘Real’) for the unit in (**a**) with the permutation-generated surrogate distribution of t-statistics (blue; box plot indicating median, interquartile range, and 5th and 95th percentiles) used to determine the significance of firing rate increase. **e**, Firing rate PSTHs of individual character-selective units (*p <* 0.01, permutation test) with mean (black line) and SE (gray shading) across all selective units. **f**, Relationship between the Memory Confidence Score (MCS) for models trained with character-selective neuron channels removed vs. the MCS for the full model. Each dot represents a single concept for a single participant. Dashed line shows equality.

## Discussion

We asked if small populations of neuronal spikes recorded within individual participants could uncover recalled memories using recordings taken during initial memory formation. While previous work has shown successful classification of on-screen characters using part of the viewing data for training and the other part to perform inference[12–14], we report a fundamentally different achievement: predicting concept identity from neuronal patterns during out-of-distribution, internally-generated reinstatement. Our deep learning method, trained on a single viewing of an audio-visual episode, successfully predicts subsequent recall of concepts using recordings from only 40-96 sites (channels) across the brain, mostly in medial temporal lobe (MTL) and frontal cortex (FC). As clinical considerations solely determine the location of recording sites, our models achieve decoding from a relatively small, nearly arbitrary complement of microelectrodes without curating brain regions or neurons with response characteristics. When we restrict our model inputs to only MTL or FC inputs, we successfully decode concepts in presleep or postsleep, respectively. Our models do not appear to take advantage of traditional single neuron concept selectivity[19], however, suggesting nonlinear interactions at the level of population neuronal spikes contain information useful to decoding of internal brain representations.

The notion that memory retrieval involves reinstatement or reactivation of the neural state active during encoding represents a central and well-supported theory in cognitive neuroscience[36], although showing reinstatement of a specific memory in a single recollection remains challenging. At the cellular level, studies using place cells in rodents demonstrate reinstatement before, during, and after sleep[37]. In humans, reactivation of MTL concepts cells was shown just prior to memory recall of their selected concepts[20] and neuronal sequences in lateral temporal cortex that occur immediately after encoding of paired words subsequently replay prior to correct cued recall[38]. These methods require identification of characteristic neurons or neural sequences to uncover internal representations, and in the latter case these neuronal sequences appear to possess a general role in representing categorical information[39]. In contrast, our work represents decoding of specific conceptual information during memory retrieval at the level of population neuronal activity without identifying neurons or sequences responsive to stimulus viewing, likely relying on dynamic interactions distributed widely over frontotemporal regions.

Systems consolidation theory posits that new memory formation depends on the hippocampus before reorganization into hippocampally-independent traces[40]. Sleep represents a critical period for this consolidation process[27], with neuroimaging studies of declarative memory showing consolidation-related changes in retrieval activity across sleep, including altered hippocampal–prefrontal connectivity[41] and a progressive shift from hippocampal to neocortical engagement[42, 43]. Our brain region–specific modeling results suggest systems consolidation can be seen at the level of population neuronal spikes (Fig. 3), as demonstrated by a striking difference between decoding of memories before sleep and after sleep. Before sleep, our “Full” model successfully decodes concept recall, while models without MTL, or without hippocampus, show little evidence of decoding. Concurrently, the “Only MTL” model, which excludes FC sites, can significantly decode concepts before sleep, while the “Only FC” model shows no evidence of decoding. Following sleep, where participants remember the episode with comparable recall performance to presleep (Fig. 1c), the “Full” model can still decode concept memories, as can the “Only FC” model, but the “Only MTL” only weakly decodes concepts. Our presleep findings are compatible with work showing reinstatement of specific memories via activity of single entorhinal-hippocampal concept neurons prior to recall vocalizations before sleep, while frontal cortex neurons with selective responses to concepts fail to do so [20]. Neurons in that study, however, were not followed to postsleep recall. Our study demonstrates reinstatement of FC activity dominates postsleep decoding, particularly considering that the “No HPC” model also fails to significantly decode concepts postsleep due to the inclusion of other regions beyond FC that possibly confuse the model (Fig. 3), suggesting that specific information in FC neurons during encoding reinstates during memory recall after sleep. While the correlation between amount of NREM sleep and FC decoding ability (Fig. 4d & Extended Data Fig. 13) supports systems consolidation occurring over one night of sleep (as shown previously for motor skill learning [44]), we do not know if an equivalent amount of wake would also demonstrate such consolidation. Future work should compare our overnight sleep results with participants that perform episodic recall before and after an equivalent period of wake, controls that have been used to show the specificity of sleep-dependent consolidation for motor learning[44, 45] and declarative memory[45–48].

The present study supports the transformation of memory representations from MTL to neocortex, highlighting the dynamic nature of the representation as long-term memories take hold. MTL and FC play central roles in memory[49] with human neuronal interareal communication between them strengthening during successful memory retrieval[50] particularly during periods of high cognitive demand[17]. Evidence suggests MTL memory independence can occur over the course of a single night of sleep[43, 51], supporting the rapid shift from MTL to FC–dependent decoding before and after sleep. Relatedly, we find FC contributes to postsleep decoding after failing to contribute to presleep decoding (Fig. 3b), as predicted by animal work showing weak FC engagement immediately after learning that reverses over days such that FC activates while the hippocampus disengages[52]. This reemergence of previously silent, memory-related neural patterns in postsleep FC may reflect activity silence [23, 53] as MTL dominates presleep reinstatement[5]. An alternative explanation could be slow learning in FC that only matches the neural patterns from viewing after overnight consolidation into existing schema[54].

Our models using only channels without concept-selective neurons show only small decreases in decoding ability (Fig. 4d) with no statistical difference from models with equivalent numbers of random channels removed (Extended Data Fig. 15). These results dovetail with our inability to decode concepts with logistic regression (Extended Data Fig. 6), as this model relies solely on tuning of neuronal spikes to concept presence during episode viewing to predict retrieval of the concept during subsequent recall. Previous results have shown that picture identity can be decoded from selective MTL neurons, but required a curated population of spike sorted neurons (19 neurons known to respond to at least one of 32 images[16]). While we find a higher proportion of selective cells (52*/*536 ≈ 10%) than previous results showing only ∼ 4% of MTL neurons respond to a set of common objects[23]), likely because we identify characters in video instead of pictures of objects, the poor performance of our logistic regression classifier indicates low neuron yields from current recording methods constrain decoding of specific memories solely from concept-selective firing rates—particularly for a one-shot experience. Confirming this limitation, recent results suggest character responsive neurons contribute little to the success of LSTM models capable of decoding character identity from intracranial neural data recorded as participants watched a TV episode[13].

Our results, therefore, suggest deep learning models take advantage of patterns in the neural code beyond single neuron tuning, which could include spike timing and nonlinear interactions between groups of neurons[33]. As transformer models have proven successful in part due to their attention mechanism, which enables, for example, language and vision decoders to capture long-range dependencies between distant words or pixels[2, 3], it is plausible that coincident neural spikes across multiple channels and timepoints similarly contribute to our decoding success. Still, the neural features underlying this decoding remain uncertain. Although training on neuronal activity during episode viewing and successfully decoding subsequent recall (and recognition, Extended Data Fig. 8) demonstrates generalization of our model across distinct memory periods, we cannot establish that the decoded signal represents an episode-specific memory trace, as concepts within a naturalistic narrative carry substantial pre-existing semantic and contextual associations. However, since microwire bundles in the participants we work with predominantly have placements in medial regions like MTL, ACC, and medial FC (73%, Extended Data Table 1) due to the nature of the electrode architecture (extending from the tip of depth probes, Fig. 1a), and these regions possess clear ties to episodic memory systems[52], we theorize that the neural patterns underlying our model success relate to memory reinstatement.

The ability to identify internal representations of specific memories from populations of unsorted neuronal spikes during verbal recall separates our model from prior brain decoding work. To this point, deep learning models trained to decode internal representations of concepts have used spike sorted single units recorded as people watch videos and have relied on dividing the viewing period into training and test[12, 13]. This previous work did not test on a true out-of-distribution period, making it difficult to know if decoding relies on perceptual responses or if it can generalize to internally-generated concepts. While scalp[18] and intracranial EEG[55] work using spectral features has shown the ability to decode semantic content of word lists during subsequent recall, these models relied on semantic similarity to classify related words on each list. Instead, our model achieves identification of specific memory retrieval during free recall of episodic content and recognition of repeated clips after training on data recorded across a single, uncurated viewing of a 42-minute episode, a relatively naturalistic use case. Decoding of memory retrieval has also been shown using multivariate pattern analysis of fMRI data, but required several viewings of a small number of short clips during training[56] and successfully decoded between only 2-3 clip identities[56] or semantic categories (e.g. face/scene) at retrieval[57, 58]. Promising new fMRI work has shown the ability to decode the gist of semantic content as single participants twice imagined telling stories they just heard[59], but required collection of over 10 hours of training data, while fMRI remains impractical for device applications. While drawbacks exist to the invasive techniques used here to measure single neuron resolution signals, our work sets a standard for what is possible for decoding specific memories using training data from one-shot learning.

Accumulating evidence suggests that deep brain stimulation can improve memory, where electrical stimulation of the entorhinal-hippocampal system[60, 61] or neocortex during awake encoding[34] or sleep consolidation[35] can enhance memory retrieval. Modulating a specific human memory, however, has remained a formidable challenge. Since our models provide a framework to decode specific memory content within individuals during reinstatement, future work timing stimulation to successfully decoded concepts has the potential to selectively enhance or mute memory for that content. Further, our ability to train single-participant models on single-shot, unsorted neural recordings enables inference of conceptual memory content in real time. The comparable performance of models using simple, low-overhead unsorted activity versus labor-intensive, spike-sorted activity (Extended Data Fig. 12) further supports the translational potential of our approach. Inference in deep learning-based architectures can be executed within tens of milliseconds on standard hardware, as demonstrated in real-time neuroprosthetic systems [31, 32], and is compatible with closed-loop stimulation applications. Methods to predict specific memory content can therefore serve as a basis for human memory neuroprosthetic devices aimed at modulating specific memories.

## Methods

### Participants

15 participants with drug-resistant epilepsy (5 females, age range 21-55 years, mean± std 35.6±12.4 years) who were implanted with intracranial depth electrodes for seizure monitoring participated in our study (12 at UCLA, 3 at Iowa). A 16th male (48 years old) participant tested at UCLA was used in Extended Data Fig. 13 only. The study protocol was approved by the Institutional Review Board (IRB) of UCLA and The University of Iowa through a reliance agreement on the UCLA IRB. All participants provided written informed consent prior to participation. We provide details on their medications and overnight sleep in Extended Data Table 4.

### Behavioral tasks

Participants viewed a 42-minute episode of 24 (Season 6, Episode 1), followed by memory tests conducted before and after an overnight sleep session (Fig. 1a). The memory tests typically consisted of three components: free recall, recognition memory, and cued free recall. P1 and P2 did not perform the cued free recall task. For recognition memory, P1 completed only the presleep test and P5 only the postsleep test; P6, P9, and P10 completed both tests, but their recognition memory data were unavailable for analysis due to recording error. In free recall, participants were asked to verbally recall all details they could remember about the episode. In the recognition memory task participants were shown 150 short clips (0.5 - 3 seconds; 75 target clips from the episode they watched, 75 foil clips from another episode in the season) in a random order and were tasked to respond whether the clip was from the episode they had watched or not. The same clips were used in both pre- and postsleep tests. In cued free recall, participants verbally answered six image-based questions each accompanied by still images of characters related to the question. These included 1) “Where does Chloe work and what upsets her about Jack?” with a solo image of Chloe, 2) “Who are these characters and how do they relate to terrorism?” with a picture of A. Amar and FBI agents, 3) “Who spoke to the president in the episode? What did they talk about?” with a solo image of the President, 4) “What was his story?” with a solo image of A. Fayed, 5) “Describe the times Jack was restrained. How did he get there or get free?” with a solo image of J. Bauer, and 6) “Where are the various places that the episode takes place? Can you describe some events from each place?” with a picture of the LA River with 3 distant figures including J. Bauer. The question text and accompanying image were removed from the screen before participants began their recall. Recall audio was recorded at 32 kHz using the Neuralynx ATLAS system.

### Neural recordings

Participants were implanted with Behnke-Fried hybrid depth electrodes (AdTech Medical) with placement determined solely by clinical criteria. Electrophysiological data were recorded using the Neuralynx ATLAS system. Intracranial EEG (iEEG; 0.1 - 500 Hz; 2 kHz sample rate) signals were recorded from macroelectrode contacts (1.5 mm wide) spaced along the probe shaft. Broadband potentials (0.1 Hz - 8 kHz; 32 kHz sample rate) were recorded from bundles of nine 40 *µ*m Platinum-Iridium microwires (8 high-impedance recording electrodes and 1 low-impedance reference wire) that were threaded through the depth electrodes.

### Electrode localization

We localized electrodes to brain regions by loading a pre-operative T1-weighted MRI (1mm isotropic) and post-operative high-resolution CT (*<*0.6mm × *<*0.6mm × 3mm) into Locate electrodes Graphical User Interface (LeGUI) software[62]. In LeGUI, the MRI and CT images were co-registered, normalized to the Montreal Neurological Institute (MNI) template, macroelectrode contacts were automatically segmented, and microelectrode contacts were manually identified from the CT scan by an expert rater. Quality control for all contact localizations was done by visual inspection during contact labeling. We segmented cortical and subcortical regions automatically using FreeSurfer [63]. Medial temporal lobe contacts, which cortical parcellation software can fail to properly identify, were automatically segmented using Automatic Segmentation of Hippocampal Subfields (ASHS) software with the ASHS ABC 3T protocol[64, 65]. Electrode contacts within the ASHS segmentations were manually labeled by visual inspection. Coordinate transformations from LeGUI to ASHS and FreeSurfer spaces was performed using FSL software[66].

### Model input data generation

#### Voltage spikes

Broadband microwire signals were notch filtered (4th order Butterworth, center frequencies from 300 to 3000 Hz in steps of 60 Hz, bandwidth 4 Hz) to reduce line noise, and high-pass filtered (4th order Butterworth, cutoff at 300 Hz) with zero-phase lag filters to isolate neural spiking events. To detect voltage spike events, we first computed the standard deviation (SD) of the filtered signal amplitude separately for each channel and task epoch (e.g., movie viewing, recall, sleep). The minimum SD across epochs for each channel was used to set a symmetric threshold at ±3 ×SD, which was then applied across all task epochs for that channel. We identified threshold-crossing events in both directions: negative-going deflections and positive-going deflections. Each threshold-crossing was considered a putative voltage spike. For negative crossings, we recorded the minimum amplitude and its corresponding time point. For positive crossings, we recorded the maximum amplitude and its time point. These values were treated as separate events to preserve the bidirectional structure of extracellular voltage signals. The resulting time series of amplitudes, one for negative and one for positive events, constituted the voltage spike amplitude trains. To reduce artifacts, we removed coincident events across channels within a microwire bundle. Specifically, if synchronous events were detected across 6 of 8 channels within a 4-ms window, we excluded these values from both the negative and positive amplitude trains. To facilitate comparability across channels and participants while preserving the relative structure of neural firing patterns, we discretized spike amplitudes into fixed bins defined by multiples of the detection threshold’s standard deviation (SD). The amplitudes were grouped into half-width intervals ranging from 3.5 to 30.5 (e.g. [3.5, 4), [4, 4.5), . . ., [30, 30.5)), and each value was replaced by the left boundary of its corresponding bin. This discretization serves to reduce sensitivity to minor amplitude fluctuations while retaining key information about relative spike intensity. Voltage spike trains were segmented into 250-ms intervals aligned with the movie episode. After excluding the end credits, the episode spanned 2,435 seconds, yielding 9,740 windowed neural segments. Each 250 ms neural signal from a single electrode was initially represented as a 2D array of shape 2 × *N*, where *N* = *f_s_* × 250*ms* = 8, 000, and the two rows encode the discretized amplitude values from negative- and positive-going threshold-crossing events, respectively. To reduce dimensionality while preserving temporal structure, we applied a non-overlapping 5-ms binning window independently to each polarity channel. Each bin contained *L* = *f_s_* × 5*ms* = 160 samples, resulting in a total of *B* = *N/L* = 50 bins per segment. Within each bin *j*, we sum the discretized amplitude values. The final processed data for each 250-ms window were structured into a tensor *w_i_*∈ R^2*×*^*^Ne×B^*, where *N_e_* is the number of electrodes, *B* = 50 is the number of temporal bins, and the first dimension indexes spike polarity, distinguishing negative- and positive-going deflections. We normalized the binned amplitudes by z-scoring them within each recording bundle, using the mean and standard deviation calculated over the entire experimental period. The normalization was performed separately for negative- and positive-polarity channels.

#### Video concept labels

We annotated the 42 minute, 24 video using custom software that calls a ChatGPT API to label frames of the video with characters. The identified concepts span three themes: characters, locations, and abstract concepts. We visualize their presence across the episode in Fig. 1c (bottom). The concepts of interest were selected based on their frequency of appearance during the episode and their frequency of mention during the recall of the participants. For “handcuff” we used the entirety of all scenes where Jack Bauer was bound; for locations like “CIA” and “White House” we used the entirety of scenes in buildings relevant to those concepts or with relevant characters on screen (e.g. the President on the phone counts as “White House”); for “sacrifice” we used the entirety of scenes Jack Bauer was being exchanged as part of a hostage negotiation or when other characters referred to Jack’s sacrifice/exchange in scenes without him. To account for temporary absences of concepts due to camera movements, scene cuts, or transitions, we filled the gaps in the labels for each concept (Fig. 1c (bottom) shows the filled labels), i.e. J. Bauer was considered onscreen as long as he was present in a given scene). Labels were aligned with the neural data for each participant using audio alignment of the episode audio track to microphone recordings taken during the experiments. Each second interval was assigned a vector as its label, where the length of the vector corresponds to the total number of concepts. Each index in the vector represents a specific concept, with 1 indicating the concept’s presence and 0 indicating its absence. This format allows multiple concepts to be assigned to the same time interval, reflecting the possibility of co-occurring concepts. To ensure sufficient training data, we divided each second into four equal parts, resulting in 250 ms annotation intervals, where each quarter-second shared the same label.

#### Recall vocalization labels

We identified the eight key concepts during vocalization of free recall and cued free recall: “White House” (includes “Washington”, “DC”, “President”, “D. Palmer” [the President’s name]), CIA (includes “FBI”, “DHS”, “CTU” [the actual name of the government organization in the show, which is typically confused for CIA], or “Chloe” [main character in CTU scenes]), “sacrifice” (includes “hostage”, “exchange”, or “trade”), “handcuff” (includes “ties”, “bindings”, “restraints”, or “chains”), J.Bauer (includes “main character” and any reference to the main agent), B.Buchanan, A.Fayed (includes “head of terrorist organization” and any reference to main bad guy setting the trap), and A.Amar (includes neighborhood “son” or “kid” in reference to his father being detained). For vocalizations of the same concept within 5 s of each other, we removed all but the first mention to avoid overlapping model predictions in our primary -4 to 0 s window. We included pronouns that refer to these concepts in our annotations when the subject referred to by the participant is clear, as we expect them to evoke similar neural reinstatement as nouns[67]. To annotate recorded participant recalls, we first implemented an automatic annotation pipeline that ran the audio files through three speech-to-text systems (Vosk [https://github.com/alphacep/vosk-api], Whisper-timestamped [https://github.com/linto-ai/whisper-timestamped], and WhisperX [https://github.com/m-bain/whisperX]) and compared generated annotations and corresponding timestamps of word vocalizations from each model. These outputs were then manually curated by a team of volunteers that fixed missing or incorrect words or timestamps using a customized version of Penn Total Recall annotation software (https://memory.psych.upenn.edu/TotalRecall). A final researcher then extracted the concepts (and their pronouns) from these corrected transcripts while listening to the audio track. Note that translation of vocalizations and accumulation into the eight categories were always finished prior to any model training and were never changed to avoid experimenter influence.

### Model architecture

Our model implementations were carried out using the Python and PyTorch frameworks. The model architecture comprises four key components: (1) a set of region embedding layers that transform neural signals from each microwire bundle into feature matrices while applying positional encodings to retain temporal information, (2) a multiregion encoder that includes region-wise self-attention layers to model intra-region relationships and cross-regional attention layers to capture interactions between different bundles, (3) a combiner module that aggregates the outputs from the multiregion encoder to produce a unified representation of collective neural activity for each 250-millisecond interval (50 time bins), and (4) a classification head comprising a single linear layer that maps the unified representation to the probabilities of the eight main concepts.

An overview of our transformer architecture is shown in Extended Data Fig. 2a. Neural signals were partitioned by microwire bundle, with each region contributing 8 electrodes to the model input. We denote the neural activity from the *R* brain regions as **x** = (*X*_1_*, . . ., X_i_, . . ., X_R_*), where each *X_i_*represents the neural signal from region *i*. The dimensions of *X_i_* vary depending on the data type. For clusterless data, each *X_i_* is represented as a tensor of shape (2, *N_e_^i^*, *B*), where *N_e_^i^* is the subset of electrodes corresponding to region *i*, and *B* is the number of time bins within the 250-ms interval. The total number of electrodes across all regions satisfies ∑*_i_*_=1_*^R^N_e_^i^*=*N_e_*. Similarly, for macro iEEG data, each *X_i_*has dimensions (1*, H_i_, T*), where *H_i_*is the subset of electrodes assigned to region *i*, each decomposed into eight frequency bands, and *T* represents the temporal dimension sampled within the 250-ms window. The total number of frequency-augmented electrodes satisfies ∑*_i_*_=1_*^R^H_i_*=*H*.

Each region-specific input *X_i_*is first processed by the region embedding layer, where it is transformed into a feature embedding augmented with positional encodings to preserve temporal information. These embeddings are processed by the multi-region encoder, which consists of two sequential mechanisms: region-wise self-attention (RSA) for intra-region dependencies and cross-region attention (CRA) for inter-region interactions. Inspired by [4], this design enables effective integration of information across regions through structured attention mechanisms. The encoded outputs are then aggregated by the combiner module, producing a unified representation of neural activity. Finally, a classification head, implemented as a single linear layer, predicts the probability that a concept is active within the given 250-ms interval.

#### Region embedding

The region embedding layer follows the patch embedding approach used in Vision Transformers (ViT) [3]. Each neural activity matrix *X_i_*is processed using a convolutional neural network (CNN), partitioning it into non-overlapping patches that serve as local feature representations. The convolution operation preserves spatial relationships while reducing dimensionality. A class token, a learnable vector widely used in ViT, is concatenated with the patch embeddings to provide a global representation of the entire region. During attention operations, this class token aggregates information from all patches, allowing the model to capture region-wide neural dynamics. The processed embeddings are then passed through positional embeddings and subsequently to the transformer layers.

#### Multi-region encoder

The multi-region encoder models both local (within-region) and distributed (cross-region) neural interactions, incorporating region-wise self-attention (RSA) and cross-region attention (CRA). The RSA layer captures intra-region dependencies, refining neural activity within each region, while the CRA layer enables inter-region interactions by integrating information across different brain regions.

#### Region-Wise Self-Attention (RSA)

The RSA layer captures interactions within the same brain region by applying self-attention to the region embeddings. Each region embedding is processed using self-attention, where queries, keys, and values are computed from the same feature matrix. The attention mechanism dynamically prioritizes relevant spatial and temporal features, refining the representation of the neural activity within each region. At the output of RSA, only the updated class token, denoted as *X_i_*, is retained, while the patch-level representations are discarded. This compressed representation of region *i* is then passed to the cross-region attention (CRA) layer, ensuring that inter-region interactions operate on compact yet informative summaries rather than the full feature maps (Extended Data Fig. 2b).

#### Cross-Region Attention (CRA)

The CRA layer models inter-region dependencies by processing the class tokens output from RSA. Unlike RSA, where attention operates within a single region, CRA computes queries from the class token of a given region, while keys and values are derived from the class tokens of all other regions. Specifically, for region *i*, the input to CRA is the updated class token *X^⎯^_i_*, while the set of keys and values is constructed by concatenating class tokens from all other regions, {*X*^⎯^*_j_* | j ≠ *i*}. This allows the model to selectively integrate inforamtion from other brain regions, capturing inter-regional interactions (Extended Data Fig. 2b). The output of CRA is an updated representation of *X^⎯^_i_*, now enriched with context from the other regions. This refined set of class tokens is then passed to the combiner module, where all regional embeddings are aggregated into a unified latent representation of neural activity.

#### Combiner

The combiner module consolidates the outputs of the cross-region attention layers to generate a final unified representation of neural activity. Prior work [4] explored both weighted summation and unweighted averaging strategies for this aggregation, reporting similar performance between the two approaches. Based on these findings, we adopt the simpler approach of averaging the regional representations to produce the final latent representation *H_t_*.

#### Further training details

We expect the overall brain state and associated neuronal activity patterns during recall of one-time viewed episodes to differ substantially from those during viewing, as recall relies on internally generated signals without concurrent sensory input. When participants view the episode, we expect two distributions of neural patterns: those responsible for perceiving the rich, continuous stream of visual and audio content and those that form memories[49]. During recall, external stimuli from the episode no longer exist, meaning the relevant reinstatement of concepts may only rescue a small component of the original audiovisual percepts[68, 69]. This difference is amplified in postsleep recall due to potential neural reorganization or electrode drift. This leads to a machine learning scenario in which training and testing data are drawn from different distributions, with only a limited shared component. Although prior work has proposed methods to address such distributional shifts through domain adaptation and neural manifold alignment[70, 71], these approaches often require densely labeled or trial-based data, which is impractical in our free-recall setting with sparse and unconstrained concept occurrences. We therefore adopt a strategy of learning the training signal as faithfully as possible and use the entire viewing period for training the model to reduce training error. During recall we expect the trained models to respond to only one of the relevant components in the neuronal activity mixture, potentially leading to an output below 0.5 even for a successfully decoded concept. This motivated our use of the Memory Confidence Score (MCS; detailed below), which compares prediction statistics before concept recall to those from matched surrogate intervals.

The model was trained using binary cross-entropy (BCE) loss for a training duration of 49 epochs with a learning rate of 1e-4, optimized using Adam and scheduled using StepLR. The transformer architecture consisted of a hidden size (feature embedding length) of 384, an intermediate (feedforward) size of 768, model depth of six RSA and CRA layers, and six attention heads (Extended Data Fig. 2). We tuned hyperparameters based on empirical performance across four participants (P3, P6, P9, P10) who exhibited rich verbal narratives during both pre- and postsleep recall. These hyperparameters included variations in model depth, hidden size, and training duration, and we selected the configuration that yielded the highest Memory Confidence Scores across these individuals. Final analysis was then performed on all participants using this set of parameters tuned to the initial four participants. Using the same four participants, we also evaluated model convergence by tracking movie-training loss and out-of-task recall MCS throughout training. Training loss progressively decreased, while recall MCS reached a stable regime around the training duration used in the paper and showed no systematic improvement with continued training (Extended Data Fig. 3). Region embeddings were extracted using CNNs with patch sizes of (1,5) for clusterless data and (1,10) for macro iEEG. To address class imbalance, we performed stratified sampling at the level of unique label combinations, ensuring proportional representation of each distinct co-occurrence pattern among the eight target concepts during training. Combinations with less than 50 seconds of total screen time were excluded, along with their corresponding neural data segments. Rather than applying cross-validation or creating held-out sets from the movie viewing data, we trained in a fully supervised manner using all available labels, prioritizing generalization to the recall phase.

### Model inference on recall data

We applied our trained models to each participants’ recall data, combining free recall and cued free recall where both were available (for P1 and P2 only free recall was available). We used the same preprocessing and voltage spike extraction methods that were applied to the training data on the microwire signals recorded during recall. As above, we partitioned data into 250 ms bins on which the model performed inference. The model output an activation between 0 and 1 for each concept independently over time (i.e. activation across concepts was not required to sum to one). The activation values indicate how strongly the neural patterns at each moment of the recall data matches the signature for the concepts that the model learned during training.

### Memory Confidence Score

To assess decoder performance during recall, we first smoothed the 250 ms binned model activations‘ for each concept using a moving average with a 1-second window. We then evaluated these smoothed activations for each concept relative to vocalization onset of that concept. For each instance in which a participant vocalized a concept, we extracted the activation signal within a 4-second window preceding the vocalization. We computed the area under the activation curve exceeding a baseline (the mean activation level for that concept across the entire recall period). The average of the areas above mean across all vocalizations of the concept was denoted as the observed activation score, *A_real_* (Fig. 2b, left, shaded region). This metric was designed to accommodate the temporal uncertainty of concept reinstatement during unconstrained free recall by integrating above-baseline activation over a relatively broad pre-vocalization window, rather than requiring a precisely timed activation peak. To evaluate significance, we performed a permutation test. For each concept, we randomly sampled the same number of 4-second windows from time points preceding unrelated words (excluding true vocalizations of the given concept, including other concept vocalizations), computed the area under the activation curve exceeding the mean activation level, and averaged them to obtain a surrogate activation score, *A_surrogate_*. This procedure was repeated 1,500 times to generate a null distribution {*A_surrogate_^(k)^*}*_k_*_=1_^1500^ (Fig. 2b, middle). The Memory Confidence Score (MCS) was defined as the percentile rank of *A_real_* within this distribution. Higher scores reflect more reliable concept-specific activations.

### Logistic regression

To evaluate whether concept decoding required the nonlinear feature transformations of the transformer architecture, we replaced the transformer architecture, including the RSA and CRA layers, with a single linear layer and flattened the input into a one-dimensional feature vector. We then trained it using the same binary cross-entropy loss with the resulting model directly mapping neural features to the eight concept outputs [72] All other training settings were identical to those used for the transformer models.

### Macrowire iEEG trained model

Macro iEEG data (2kHz sampling) were bipolar referenced before spectral decomposition using a Morlet wavelet transform, extracting the instantaneous powers of eight logarithmically spaced frequency bands ranging from 3 to 180 Hz [34]. As with the microwire model input, each training data point was represented as a tensor *w_i_*’ ε R^1×*H*×*T*^. Here, *H* = *N_e_*’*8 corresponds to the appended number of re-referenced electrode pairs *N_e_*’, each decomposed into eight frequency bands, and *T* corresponds to the temporal dimension derived from the 250-ms window sampled at *f_s_*’. The model architecture, training procedure, and MCS evaluation metrics remained as described above.

### Circular shift permutation test

To evaluate whether the decoder performance reflected meaningful structure in the data rather than random associations, we performed a circular shift permutation test (n=100 permutations) that disrupts label alignment while preserving the temporal continuity of both the neural input data and movie label data. Specifically, we displaced the neural data along the time axis by the same magnitude across all channels, with any data that extended beyond the end of the time axis wrapping back to the beginning. Within a given permutation the neural data for each participant was shifted by the same magnitude. This preserves the internal temporal structure within each 250-ms bin, as well as the vast majority of the structure across bins, but breaks the alignment with the movie-derived labels. This method serves as a conservative control, particularly given the structured nature of the movie stimulus, which includes correlated and repeating events. For each of the shifted datasets we trained transformer models for each participant, performed inference on the unaltered recall data, and calculated a Memory Confidence Score (MCS) for each concept using the same procedure as described above. We performed a Wilcoxon signed-rank test against 50% for the pooled MCSs across participants in each permutation to generate a surrogate distribution of 100 Z-statistics for presleep recall and 100 Z-statistics for postsleep recall.

### Statistics

We perform statistics on grouped MCS across all concepts and participants to determine if our models can decode memories from out-of-sample recall data above chance level. When comparing the distribution of MCS for a given model, we use a Wilcoxon signed-rank test to assess statistical significance against 50%. We also provide Cohen’s D in each case to measure effect size. When comparing models directly, we use a two-sided Wilcoxon signed-rank test between matched participant-concept observations. For analyses involving repeated observations, we use linear mixed-effects models implemented with statsmodels.MixedLM and fitted using restricted maximum likelihood. Random intercepts account for the relevant clustering of observations within participants, concepts or concept pairs, and random masks, as specified for each analysis. Fixed effects are reported as *β* ± SE, with two-sided Wald-test *P* values. To assess relationships between concept-specific recall counts and MCS, we use ordinary least-squares regression with concept identity included as a fixed effect and participant-clustered standard errors. Recall counts are natural-log transformed, while overnight changes in recall count are transformed as sign(Δ*N*) log(1 + |Δ*N* |). Regression coefficients are reported as *β* ± SE with two-sided *P* values. We provide a summary table of all tests in Extended Data Table 2.

### Sleep scoring

Sleep scoring follows the automated procedure of [35]. We identified NREM epochs in a clean intracranial EEG channel by estimating slow-wave and spindle-band power in 30 s windows and clustering with a two-component Gaussian mixture model to separate NREM from desynchronized REM/wake epochs.

### Character selective units

#### Spike sorting and unit analysis

We used WaveClus3 [73] for automated spike detection and sorting. We then manually curated the automated sorting output by evaluating the waveform shape, inter-spike interval violations, and firing consistency across recording sessions. Units with a firing rate across the video viewing period of *<* 0.5 spk/sec were excluded from further analysis. Z-scored firing rates were calculated using the mean and standard deviation of the binned firing rate (100 ms bins) across the full episode.

#### Character labeling and mapping to concepts

To annotate the on-screen presence of 10 characters in the 24 episode at frame-level resolution (29.97 frames per second) we used custom Python software which queried the ChatGPT API with images from the first frame of each camera cut. The model returned characters present in each image, and these labels were then applied to all frames within the shot. Character presence was determined solely by their visual appearance in each frame. Frame-level labels were manually curated. The 10 characters we labeled were unique to the concepts used for our decoding. Some decoded concepts mapped directly to characters (A.Amar, A.Fayed, J.Bauer) whereas for other concepts we assigned multiple characters (K.Hayes, T.Lennox, W.Palmer for ‘WhiteHouse’; C.OBrian, N.Yassir, M.OBrian, M.Pressman for ‘CIA’).

#### Character selectivity analysis

For a character appearance to be included in this analysis we required that the character appeared alone onscreen for at least 1 second. For each unit we performed a paired t-test comparing the firing rate in a 1 second pre-onset window (-1 s to 0 s relative to character onset) to 1 second post-onset (0 s to 1 s) for all appearances of a given character. We then compared the observed t-statistic to a surrogate distribution of t-statistics generated by randomly flipping the sign of the pre-to-post onset differences of each individual appearance (n = 1000 permutations). A unit was deemed selective to a character if (1) the unit fired at least 1 spike in more than 30% of character appearance post-onset windows, (2) the observed t-statistic exceeded at least 99% of the permuted distribution (*p <* 0.01, one-tailed), and (3) the unit only significantly responded to one character. Running the above analysis with a permutation significance threshold of *p <* 0.05 did not significantly alter the results. We also defined ‘Responsive’ units that adhered to constraints (1) and (2) above, but could respond to more than one character (see Extended Data Fig. 14b-d).

## Data and code availability

In line with protocols approved by the UCLA Institutional Review Board for protection of patient confidentiality data supporting the findings of this study is available from the corresponding authors upon reasonable request for use in collaborative research. We provide code to run models at: https://github.com/roychowdhuryresearch/human-memory-decoder.git

## Acknowledgments

We thank Christopher Garcia for participant data collection at the University of Iowa, Aditya Datta, Andreina Hampton, and Anthony Rangel for participant data collection at UCLA, James Bruska and Abdullah Akbar for technical assistance with annotating vocalizations (J.B.,A.A.) and frame-level character labeling (A.A.), Ben Falkenburg for contributions to spike sorting, and Aatmi Mehta, Arya Bhalla, Brandon Thich, Louisa Zhao, Lourin Abdalla, Mika Geiger, Rachel Jordan, Suheon Park, Teshinee Kukamjad, and Yuna Bi for contributing to recall annotations. This work was supported by National Institute of Neurological Disorders and Stroke grants R01NS084017 and U01NS123128).

## Author information

### Author contributions

Conceptualization: Z.A., V.R. and I.F. Supervision: J.S., V.R. and I.F. Formal analysis: Y.D., S.D., J.S., C.D. and Y.Z. Methodology: Y.D., S.D., J.S., Z.A., V.R. and I.F. Data collection: S.D., J.S., C.C., J.B., A.R. and K.N. Surgeries: C.C., H.K., M.H. and I.F. Funding acquisition: M.H., V.R. and I.F. Manuscript writing: Y.D., S.D., J.S., V.R. and I.F.

### Corresponding authors

Correspondence to Vwani P. Roychowdhury or Itzhak Fried.

## Extended data

**Extended Data Fig. 1.**
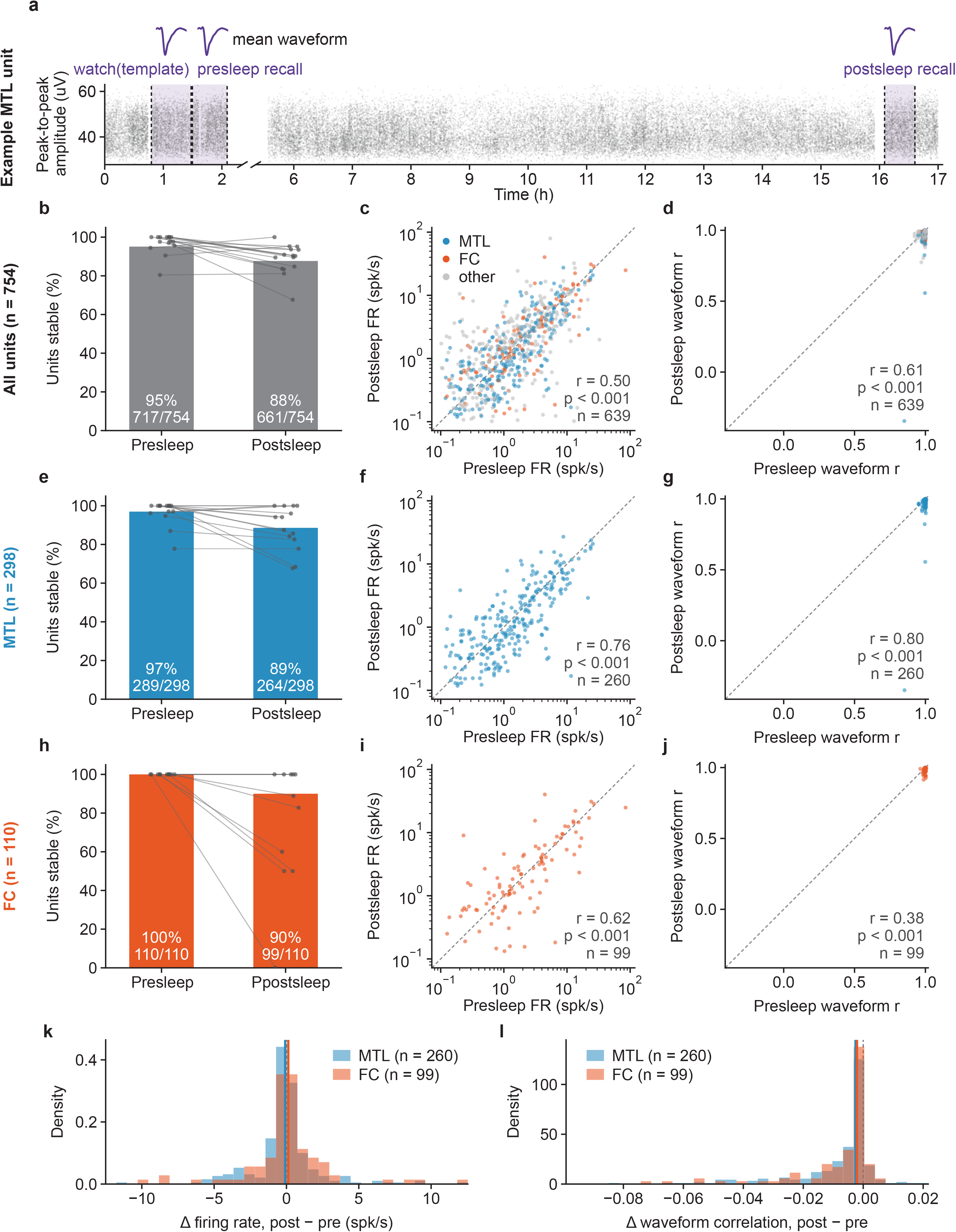
Unit stability across the full experimental series. **a**, Example curated unit over the full experimental series for one participant. Each point represents the peak-to-peak amplitude of one spike. Purple shaded epochs indicate the movie watch period, presleep recall, and postsleep recall. Above each epoch is the mean waveform of spikes recorded during that epoch. Units with a firing rate *>* 0.1 spk/sec during the movie watch period were included in the stability analysis. **b**, Percentage of units firing at *>* 0.1 spk/sec during each recall period across all 15 participants; the denominator is the number of units active (*≥* 0.1 spk/sec) during the movie watch period. Units that maintain a firing rate *>* 0.1 spk/sec during both pre- and postsleep recall are considered “stable”. **c**, Firing rates of stable units during pre- and postsleep recall. **d**, Pearson correlation between each recall-period mean waveform and the watch-period mean waveform for stable units. **e–g**, As in **b–d**, for MTL units only. **h–j**, As in **b–d**, for FC units only. **k**, Postsleep *−* presleep change in firing rate; median *−*0.089 spk/sec (MTL, *n* = 260) versus +0.126 spk/sec (FC, *n* = 99), Wilcoxon rank-sum, *p* = 0.120. **l**, Postsleep *−* presleep change in waveform correlation; median *−*0.003 (MTL, *n* = 260) versus *−*0.002 (FC, *n* = 99), Wilcoxon rank-sum, *p* = 0.708.

**Extended Data Fig. 2.**
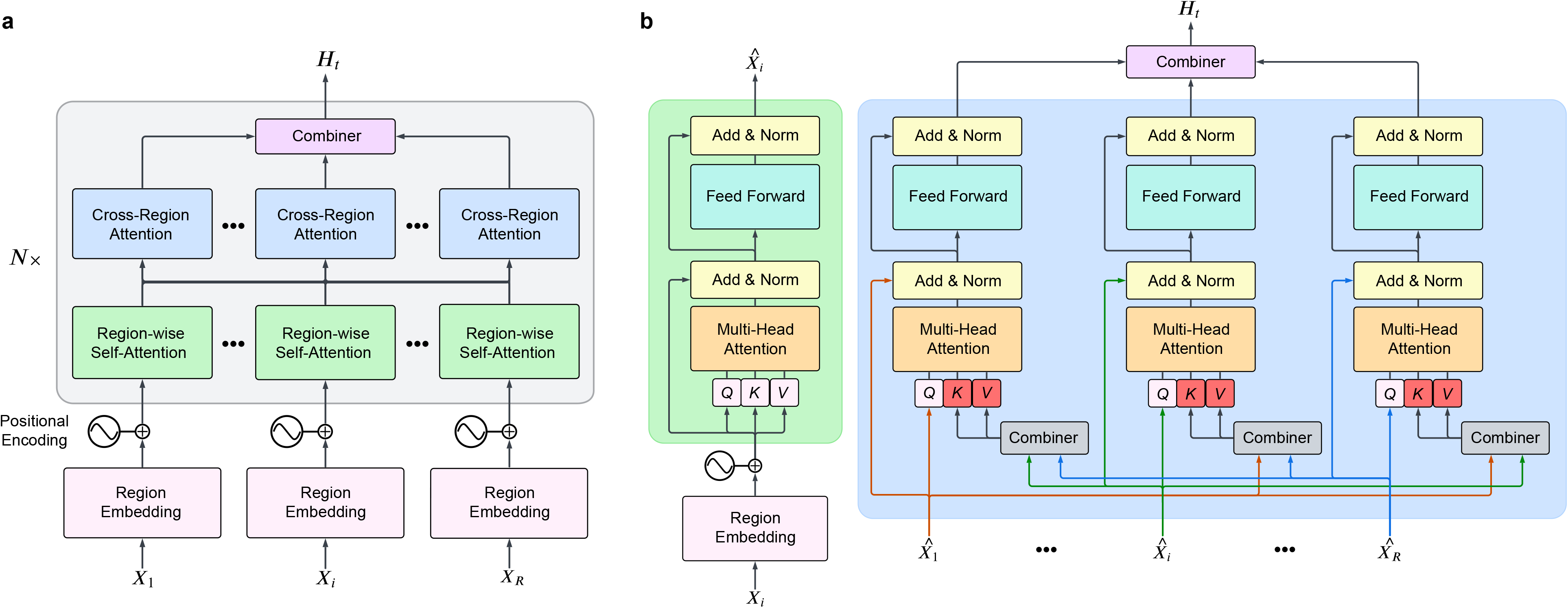
Overview of the multi-region transformer architecture. **a**, We process neural signals (*X*_1_*, . . ., X_i_, . . ., X_R_*) from distinct brain regions through region-specific embedding layers, followed by region-wise self-attention and cross-region attention layers, and finally integrate by a combiner module to produce a unified representation of neural activity *Ht.* The classification head predicts the probabilities of eight target concepts for each 250-ms interval. **b**, Detailed structure of the Region-wise Self-Attention (RSA) and Cross-Region Attention (CRA) layers. The RSA layer captures intraregion dependencies by attending to relationships within the same region, including interactions with the class token. The CRA layer processes outputs from RSA layers, enabling interactions across brain regions by aggregating information from all class tokens. Arrows illustrate the data flow within and across regions.

**Extended Data Fig. 3.**
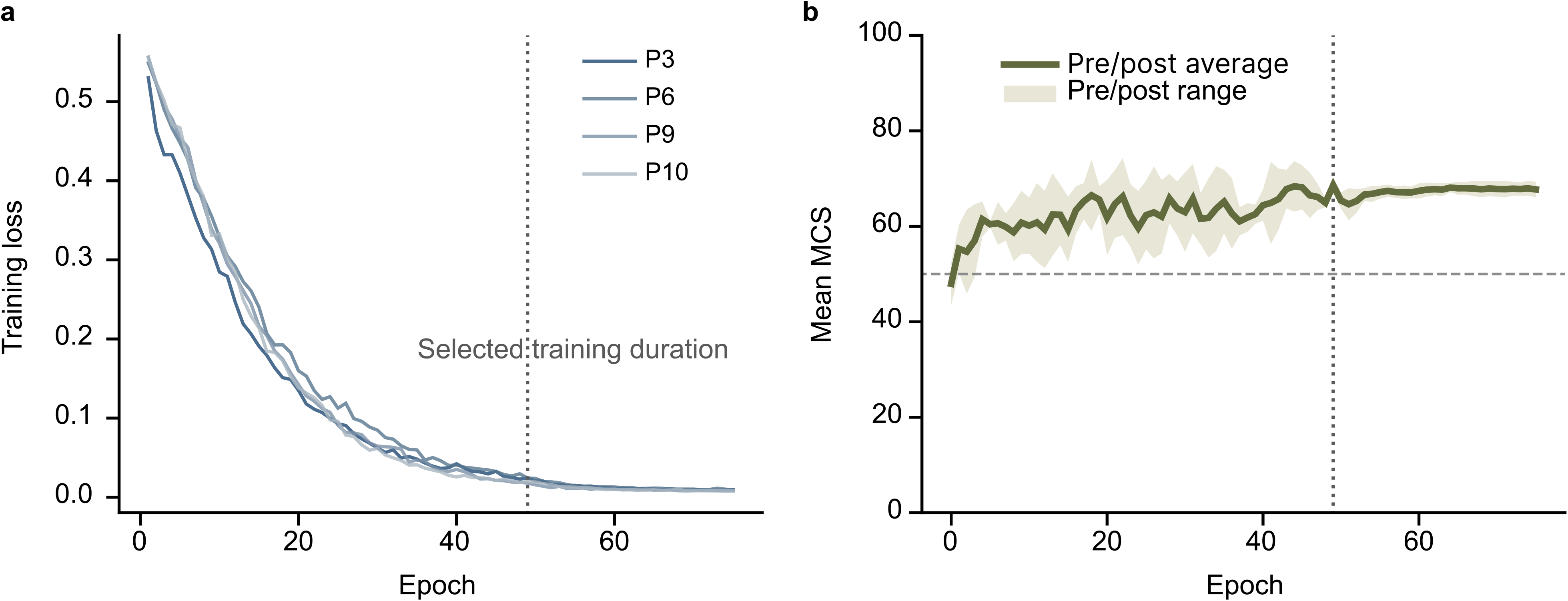
Convergence of movie-trained decoding models. **a**, Training loss across 75 epochs for the four participants used to select model architecture and training duration (P3, P6, P9, and P10). **b**, Out-of-distribution recall decoding performance, quantified using MCS. We average across eligible concepts within each participant and then across participants. The solid line shows the average of the presleep and postsleep mean MCS, and the shaded envelope spans the presleep and postsleep means at each epoch. The horizontal dashed line indicates chance-level MCS (50). The vertical dotted lines in a and b indicate the final training duration used in all analyses (epoch 49).

**Extended Data Fig. 4.**
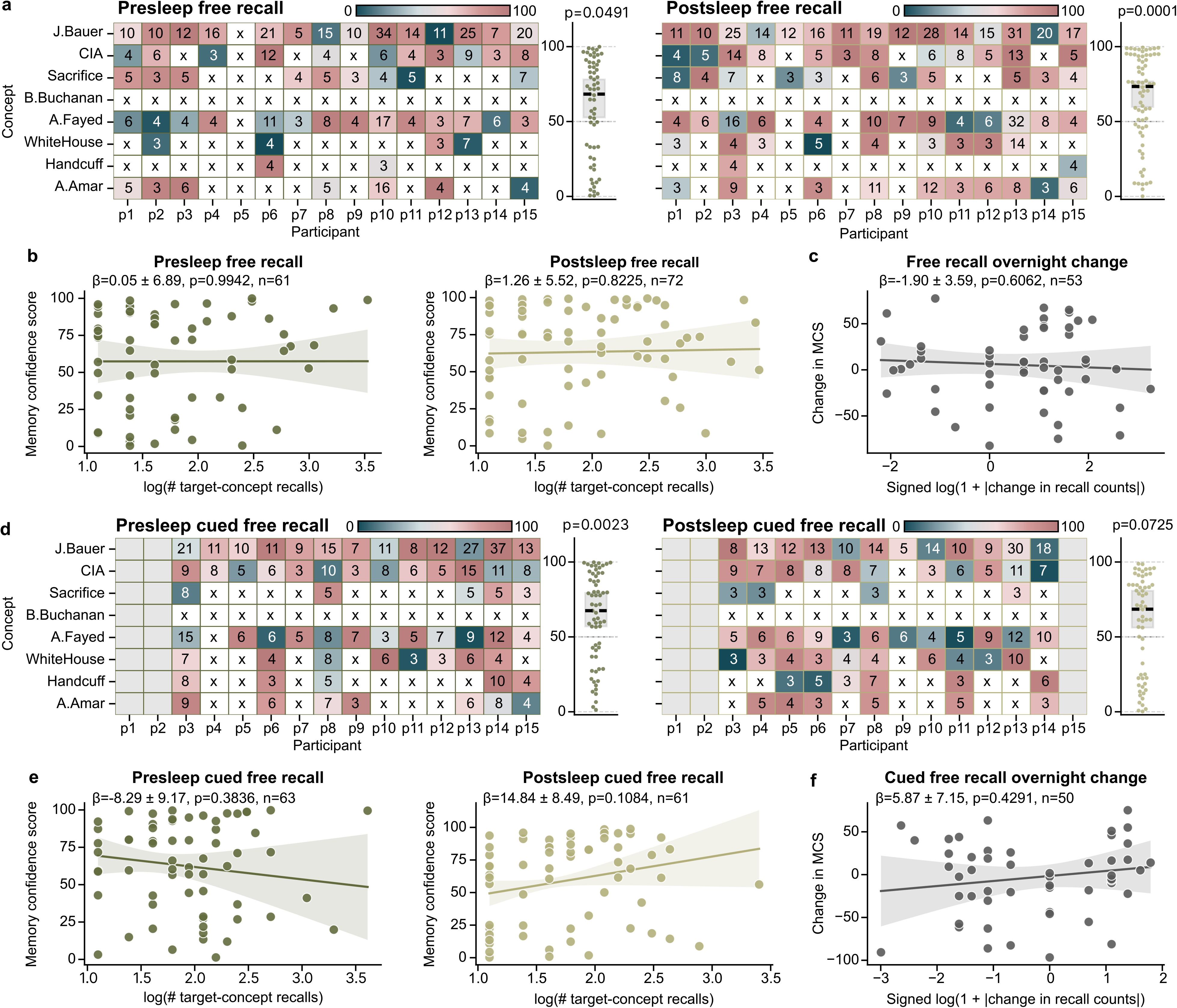
Decoding and behavioral performance during free and cued free recall. **a**, MCS during presleep and postsleep free recall. Same conventions as Fig. 2. **b**, Relationship between log-transformed target-concept vocalization counts and MCS during presleep and postsleep free recall. Each dot represents one concept within one participant. **c**, Relationship between the overnight change in recall count and the corresponding change in MCS for participant-concept pairs available in both recall periods. Recall-count changes were transformed as sign(Δ*N*) log(1 + |Δ*N* |), with changes calculated as postsleep minus presleep. **d–f**, Corresponding analyses for cued free recall; gray columns indicate unavailable task sessions. In **b,c,e,f**, lines and shaded regions show fitted relationships and 95% confidence intervals from linear regression models accounting for concept identity and clustering within participants.

**Extended Data Fig. 5.**
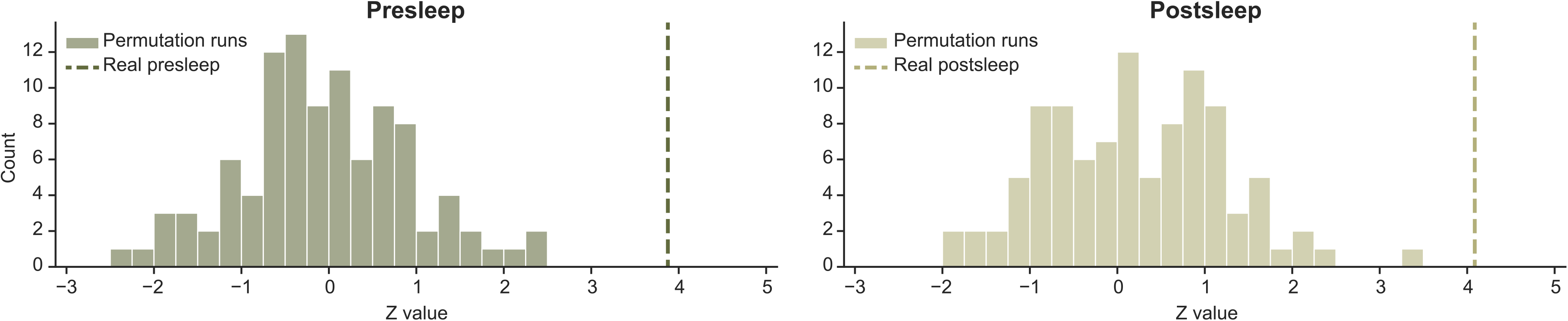
Results of circular shift permutation tests. **a,** Histogram of Z statistics from Wilcoxon signed-rank tests for each of the 100 models with randomly circularly-shifted neural data in reference to the episode annotations (Fig. 1e) for presleep memory tests. We use the same procedure as shown in Fig. 2a-b to calculate MCS and compare to 50% across participants for each model. Dashed line shows the Z statistic for correctly-aligned data *Z_real_* = 3.88 (100th percentile). **b,** Same for postsleep recall. Dashed line shows Z statistic for correctly-aligned data *Z_real_*= 4.07 (100th percentile).

**Extended Data Fig. 6.**
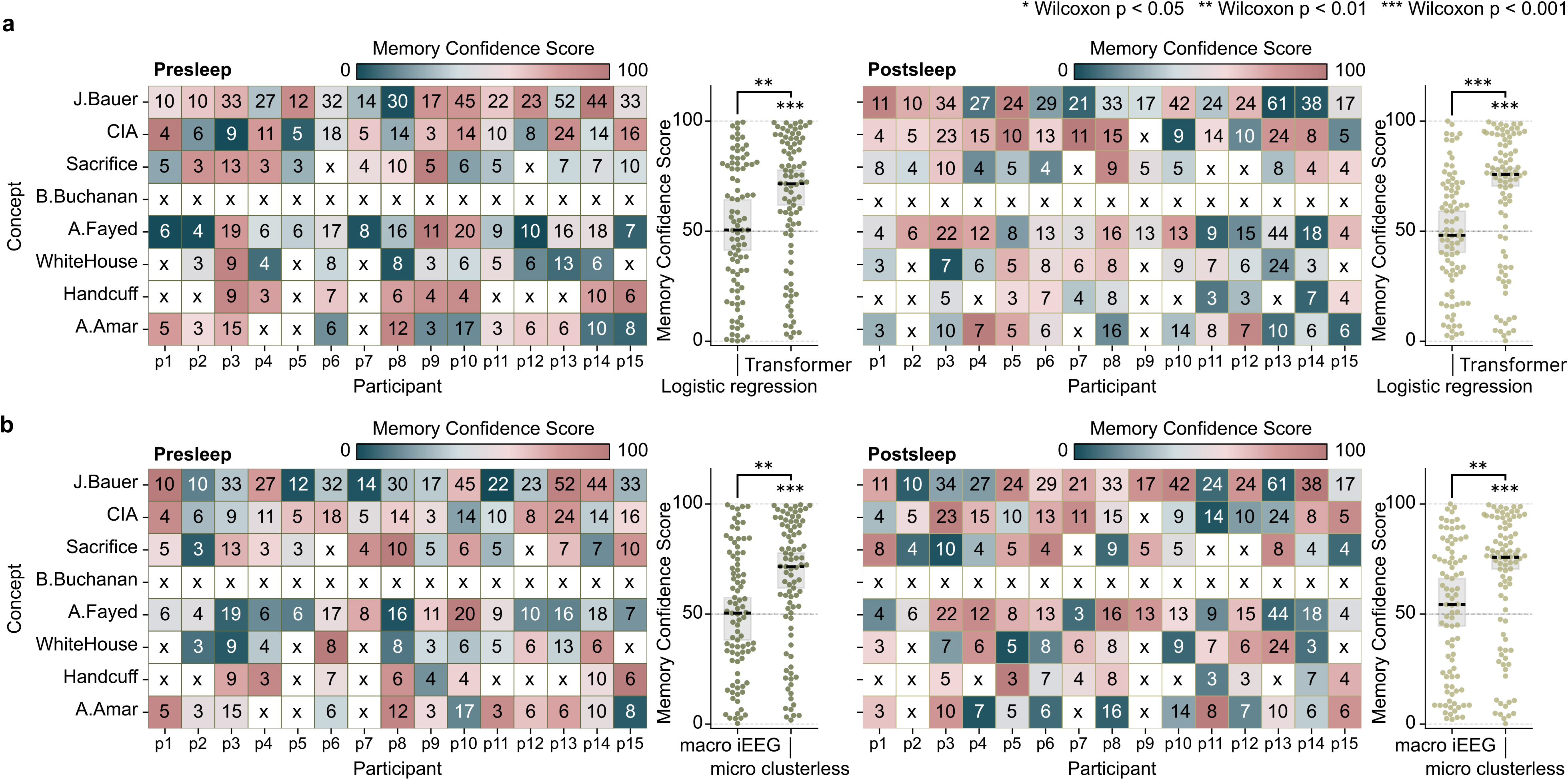
Control models fail to show decoding. **a,** For both pre- (left) and postsleep (right) memory tests we use a separate multi-label logistic regression model (Methods) for each participant to decode recall of the same eight concepts and test the performance by calculating MCS as in Fig. 2a-b. Our transformer model performs significantly better than logistic regression for both pre- and postsleep. **b,** For both pre- (left) and postsleep (right) memory tests we run our transformer model on whole-brain, bipolar-referenced macrowire iEEG broadband oscillatory power inputs and test them using the same MCS procedure as in Fig. 2a-b. These data do not reflect neuron-level spikes in voltages but instead aggregate voltages across hundreds of thousands of neurons for each bipolar channel. Our transformer model with microwire inputs performs significantly better than the model with macrowire inputs for post pre- and postsleep. Statistical tests on individual distributions indicate Wilcoxon signed-rank tests against chance (50%), and tests between paired distributions (shown by brackets) represent Wilcoxon signed-rank tests. \**p <* 0.05, \*\**p <* 0.01, \*\*\**p <* 0.001.

**Extended Data Fig. 7.**
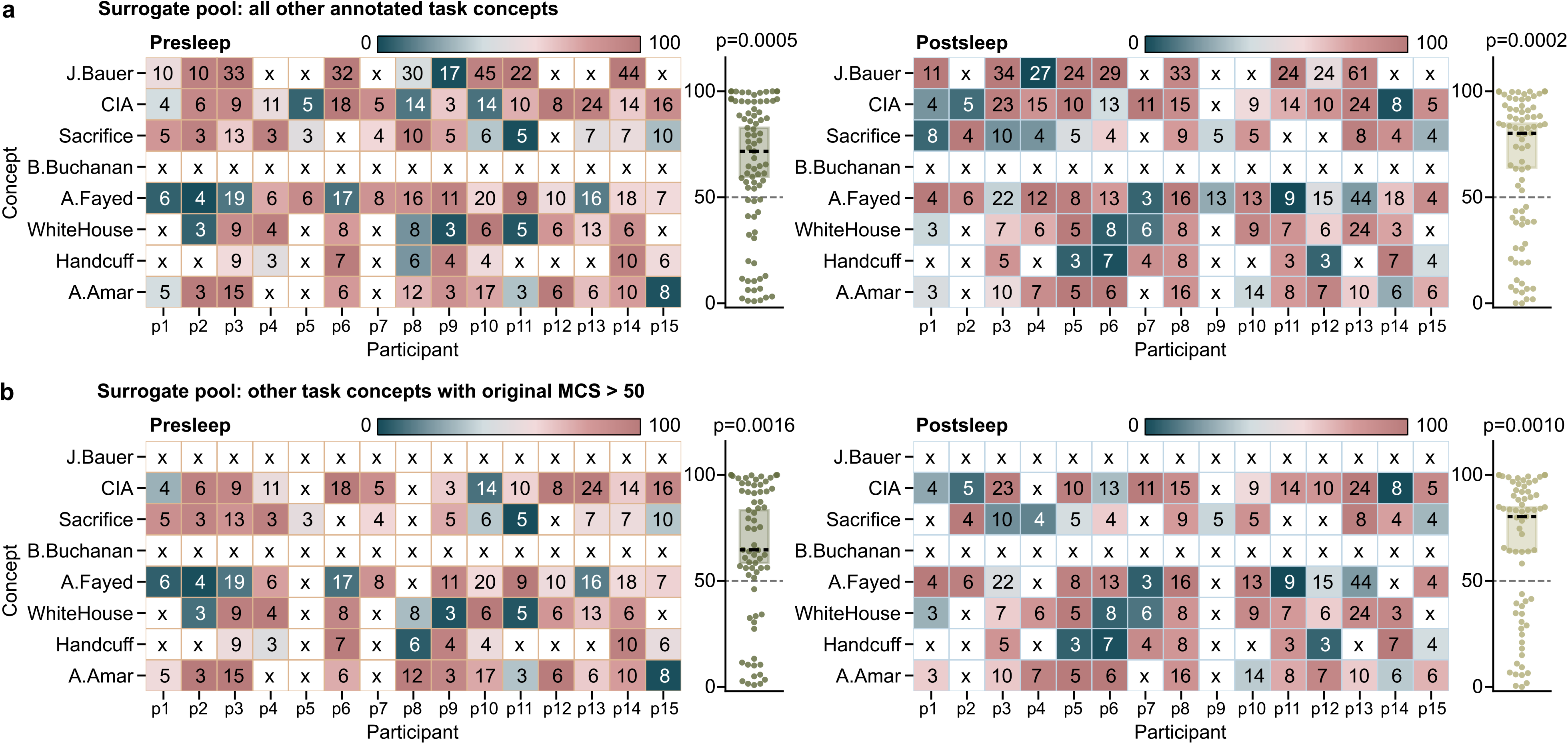
Memory Confidence Score (MCS) using surrogate vocalizations from other annotated concepts confirms significant decoding. **a**, MCS obtained when the surrogate pool comprised vocalizations from the seven other annotated task concepts within the same participant and recall period. For each permutation we randomly sample a surrogate pool as in Fig. 2b but always resample from this same concept vocalization pool. **b**, More stringent analysis restricting the surrogate pool to only the subset of the other seven concepts with an original MCS (Fig. 2d) above 50. In both analyses, we required the surrogate pool to contain at least 1.1x as many valid vocalizations as the target set. Crosses indicate insufficient target vocalizations or surrogate windows. All other conventions same as Fig. 2d.

**Extended Data Fig. 8.**
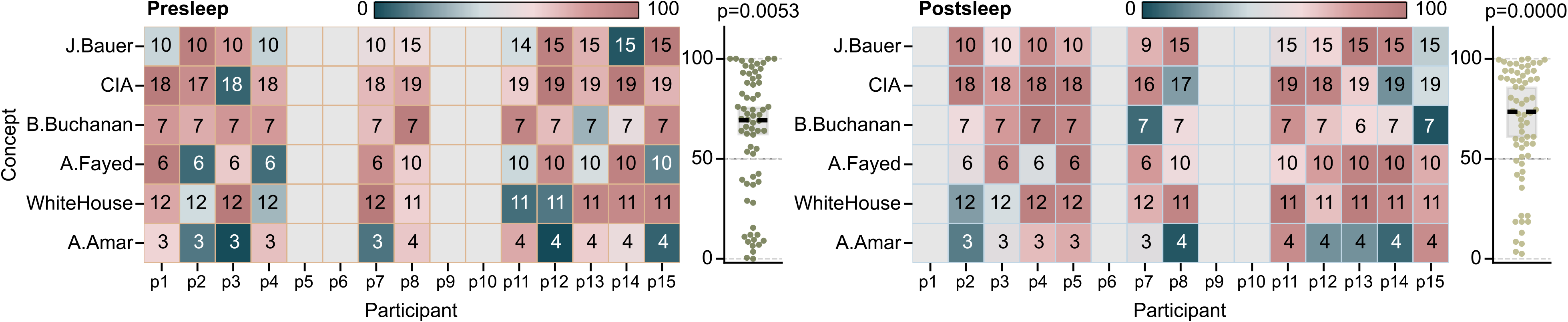
Concept decoding during recognition clip viewing. We applied frozen movie-trained models—without additional training or fine-tuning—to neural recordings taken during the half of trials with repeated clip presentations during the recognition memory task. Recognition MCS quantifies whether decoder activation within 2 s windows centered on annotated clip midpoints is higher for clips containing a concept than for concept-absent clips (which we randomly sample to create the surrogates). Numbers in each cell indicate the number of concept-present clips and empty cells indicate unavailable sessions. Same conventions as Fig. 2d.

**Extended Data Fig. 9.**
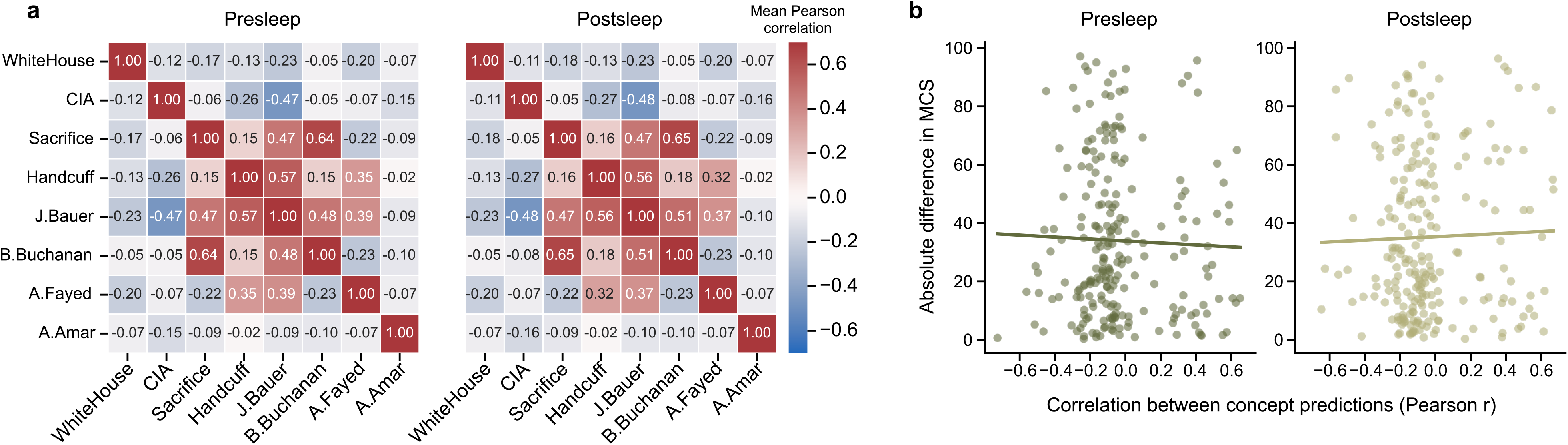
Correlations between continuous concept predictions and concept decoding performance. **a**, Pairwise Pearson correlations between continuous concept-prediction time series, calculated separately within each participant’s presleep and postsleep recall sessions and aggregated across participants using Fisher’s *z* transformation. **b**, Relationship between the concept-pair correlations in **a** and the absolute difference in MCS for each concept pair within each participant, separately for presleep and postsleep. Lines show fixed effect estimates from mixed effects models with random intercepts for participant and concept pair. No significant relationship was observed during presleep (*β* = *−*3.35 *±* 7.12, *p* = 0.638, *n* = 227 participant–concept-pair observations) or postsleep (*β* = 2.98 *±* 7.10, *p* = 0.675, *n* = 229); estimates are *β ±* SE.

**Extended Data Fig. 10.**
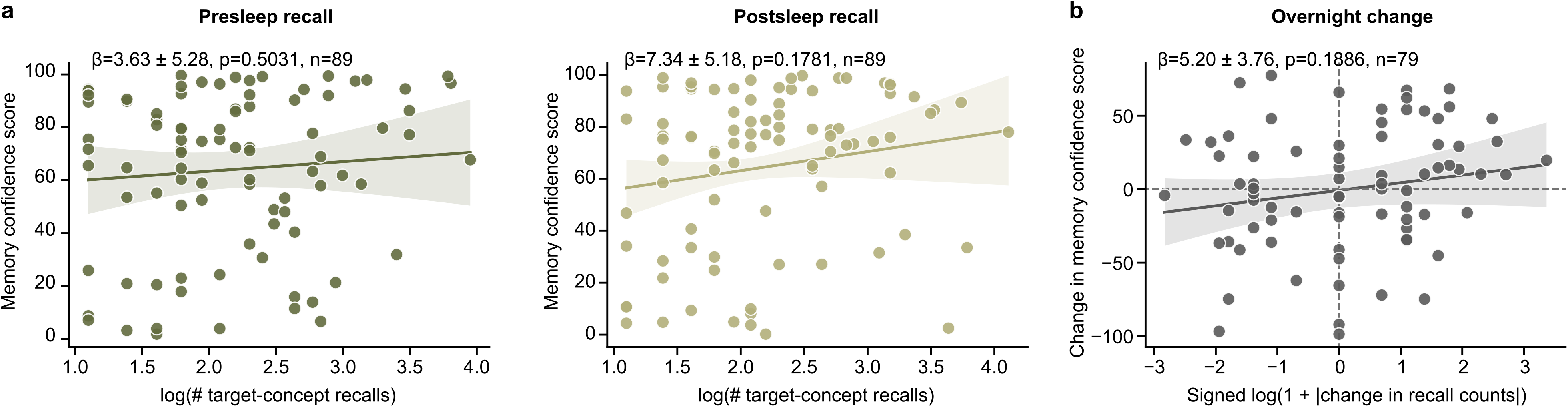
Association between concept-specific recall frequency and memory decoding. **a**, Relationship between the log-transformed number of target-concept vocalizations and MCS during presleep and postsleep recall. Each dot represents one concept within one participant. **b**, Relationship between the overnight change in recall count and the corresponding change in MCS for participant-concept pairs available in both recall periods. Changes were calculated as postsleep minus presleep, with recall-count changes transformed as sign(Δ*N*) log(1 + |Δ*N* |) to account for the log of the change. Lines and shaded regions show fitted relationships and 95% confidence intervals from mixed effects models accounting for concept identity as fixed effects and participants as random effects.

**Extended Data Fig. 11.**
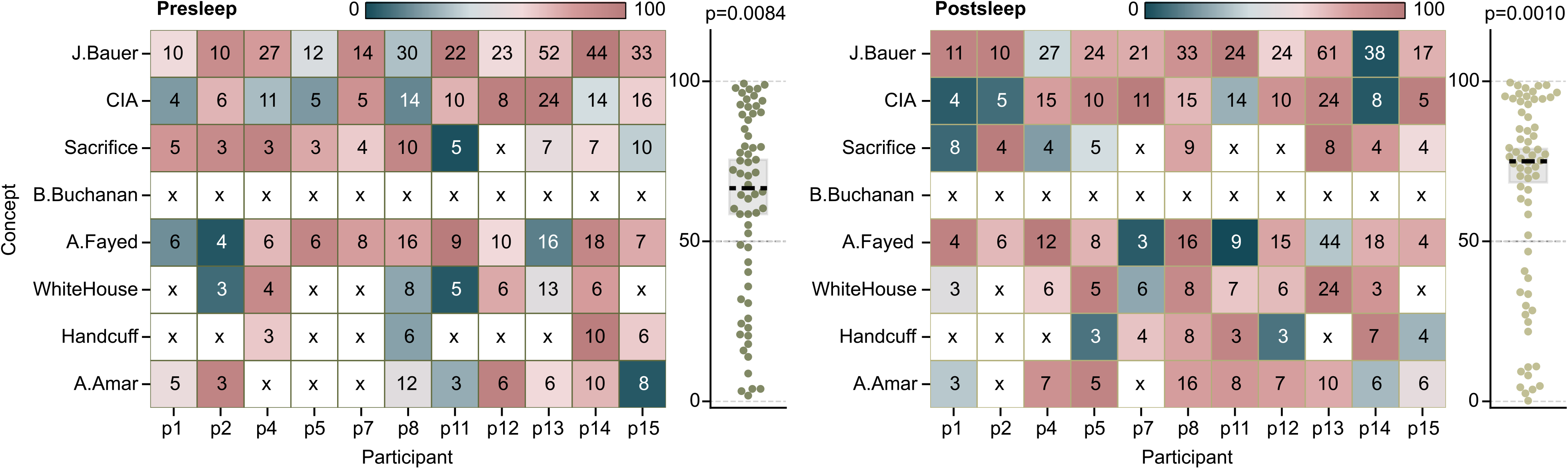
Memory decoding in participants not used for hyperparameter tuning. MCS values from the full models are shown for the 11 participants not used to select the model architecture or training duration, meaning we exclude P3, P6, P9 and P10. Same conventions as Fig. 2d.

**Extended Data Fig. 12.**
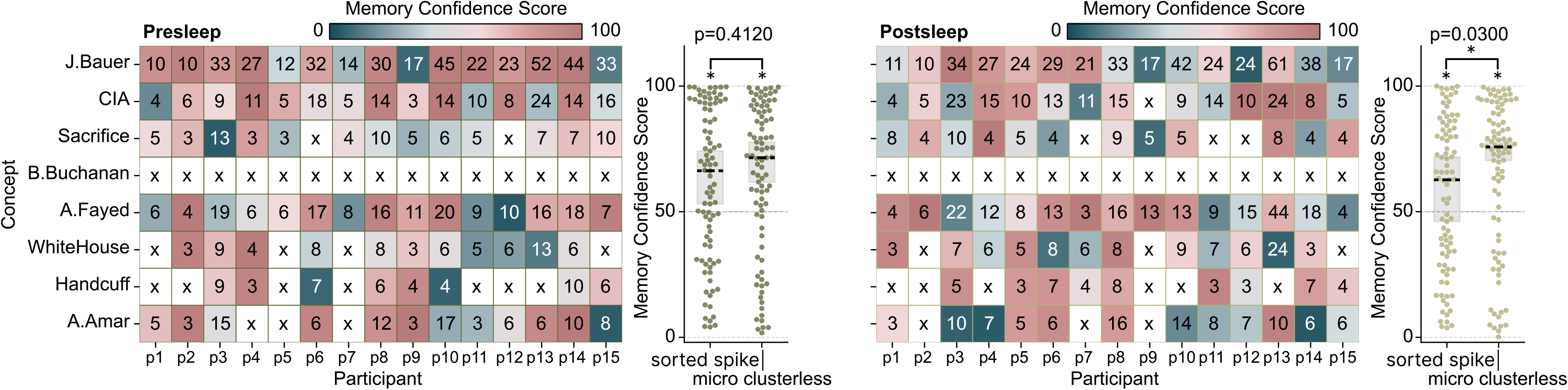
Concept decoding using sorted spike activity. MCS for full models trained using manually sorted spikes during presleep (left) and postsleep (right) recall. Numbers indicate target-concept vocalizations; “x” indicates fewer than three vocalizations. MCS was above chance during presleep (*p* = 0.0012) and postsleep (*p* = 0.0408, *n* = 89; Wilcoxon signed-rank tests). Comparisons to Full model: presleep, *Z* = 0.820*, p* = 0.4120*, D* = 0.068; postsleep: *Z* = 2.170*, p* = 0.0300*, D* = 0.214 (paired Wilcoxon signed-rank tests and paired-samples Cohen’s *D*).

**Extended Data Fig. 13.**
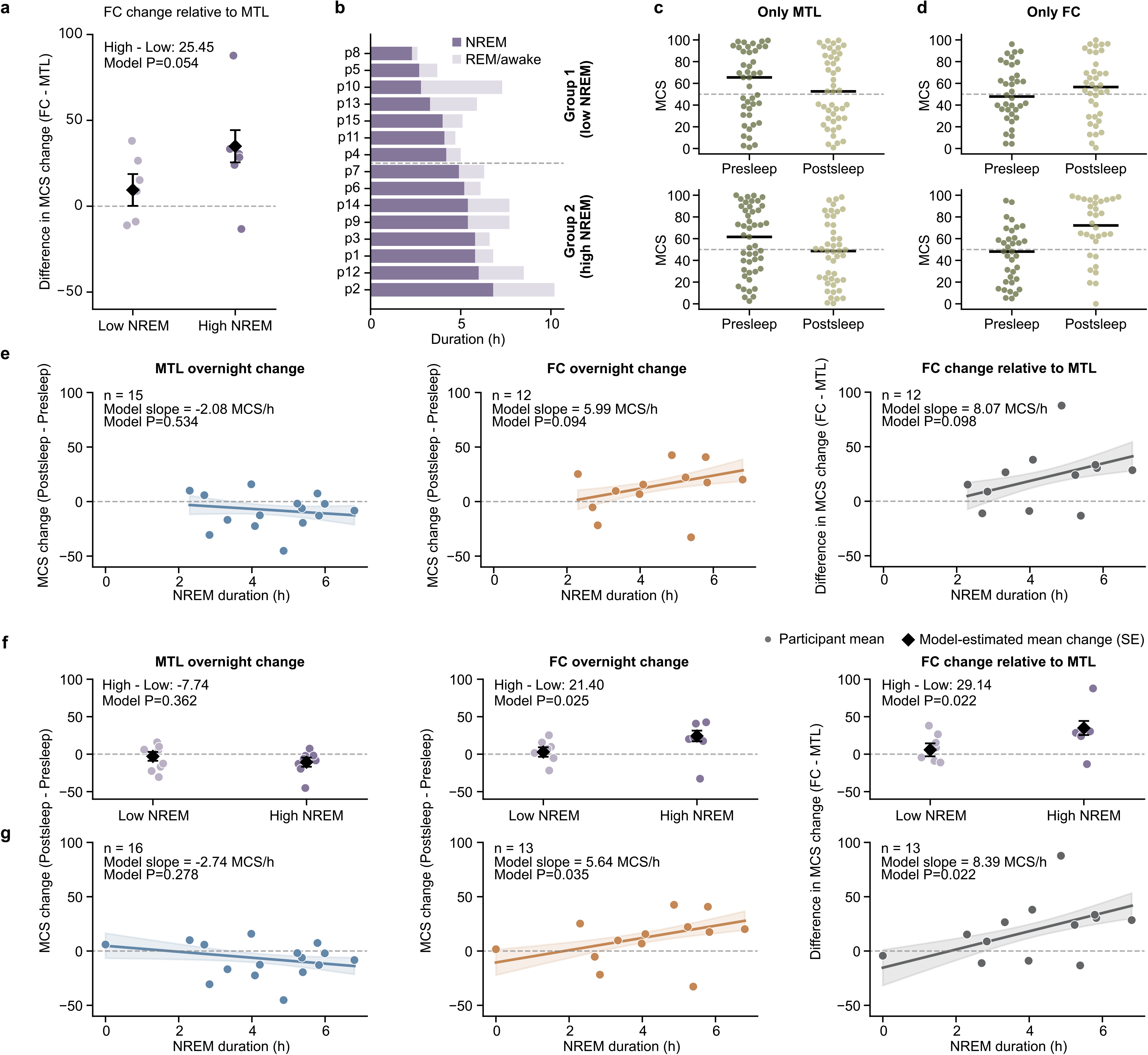
NREM sleep relates to regional changes in memory decoding. **a**, Difference in MCS change (postsleep - presleep) between the Only FC and Only MTL models for high vs. low NREM groups among P1-P15. **b**, Total recorded overnight duration for P1-P15, partitioned into NREM and REM/awake time and ordered by NREM duration. **c,d**, Presleep and postsleep MCS distributions for the Only MTL (**c**) and Only FC (**d**) models for low (top) vs. high (bottom) NREM groups. Dots represent participant-concept observations, black horizontal lines indicate medians, and dashed lines indicate chance-level MCS (50). **e**, Relationships between NREM duration and participant-level MCS change for the Only MTL model, Only FC model, and their difference (FC - MTL) among P1-P15. **f,g**, Corresponding high vs. low NREM (**f**) and continuous (**g**) analyses after including P16, a participant that performed the two memory tests 8 hours apart on the same day with 0 h NREM in between. In **a,f**, colored dots represent participant means and black diamonds and error bars show model-estimated means *±* s.e. In **e,g**, lines and shaded regions show model-estimated relationships *±* s.e. *P* values were obtained from mixed effects models with participant-level random intercepts.

**Extended Data Fig. 14.**
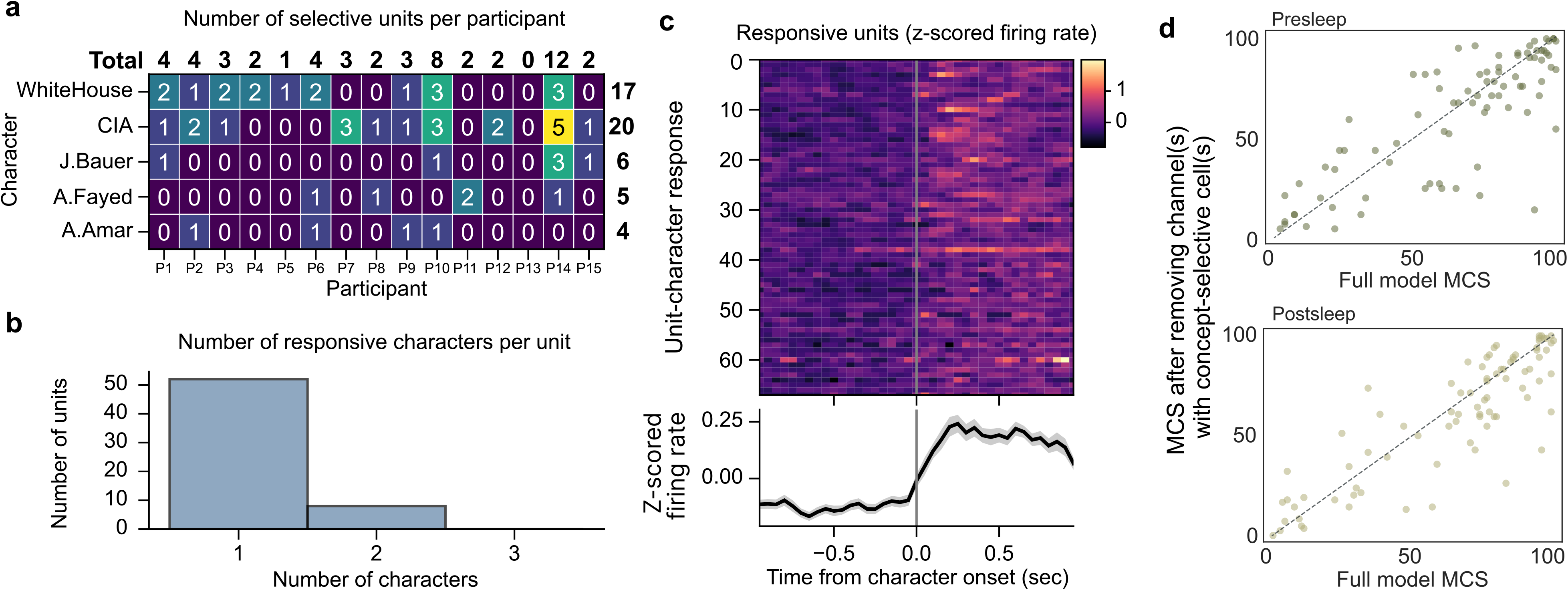
Multi-character–responsive units cannot explain decoding performance. **a**, Significant responsive units by participant for the 52 neurons selective to only one character (*p <* 0.01, permutation t-test). **b**, Eight additional units respond significantly to more than one character. **c**, Z-score firing rate PSTHs of significant character responses, including units with responses to *>* 1 character (52 single and 8 double for 68 rows) (*p <* 0.01, permutation test) with mean (black line) and SE (gray shading) across all units at bottom. Z-score calculation as in Fig. 4. **d**, Relationship between full-model MCS and MCS after removal of channels containing character-responsive units, including units responsive to one or multiple characters. Mean participant-level changes in MCS (channel removal minus full model) were -2.52 during presleep recall (*Z* = *−*0.85*, p* = 0.397*, D* = *−*0.291) and -2.98 during postsleep recall (= *−*1.54*, p* = 0.124*, D* = *−*0.514; paired Wilcoxon signed-rank tests and paired-samples Cohen’s *D*; *n* = 14 participants). Same conventions as Fig. 4f.

**Extended Data Fig. 15.**
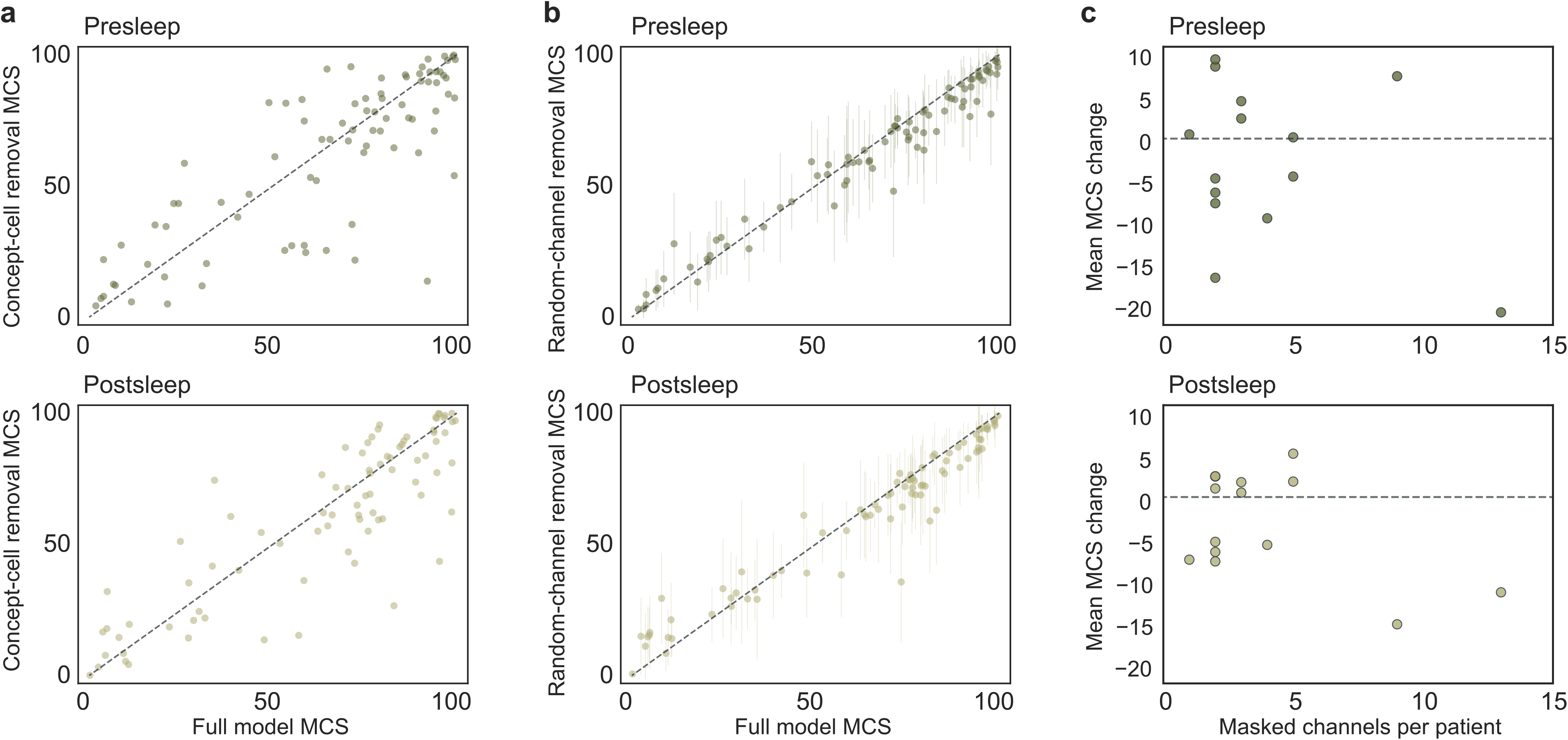
Removing channels containing significant character responses does not significantly impact memory decoding. **a**, Concept-level MCS obtained using the full model vs. MCS after masking channels containing units with significant character responses, including units responsive to more than one character (same data as Extended Data Fig. 14f shown for comparison convenience). **b**, Matched random-channel control. For each participant, the same number of channels without significantly character-responsive units was removed, with bundle occupancy matched to the character-responsive channel mask. Points show mean MCS across 60 random masks and error bars indicate the standard deviation across masks. Note that the tighter distributions of MCS for the random-channel removal than **a** are expected due to the averaging across many model runs in **b**. **c**, Participant-level mean change in MCS after channel removal as in **a**, plotted against the number of masked channels. Dashed lines indicate no change.

**Table S1.**
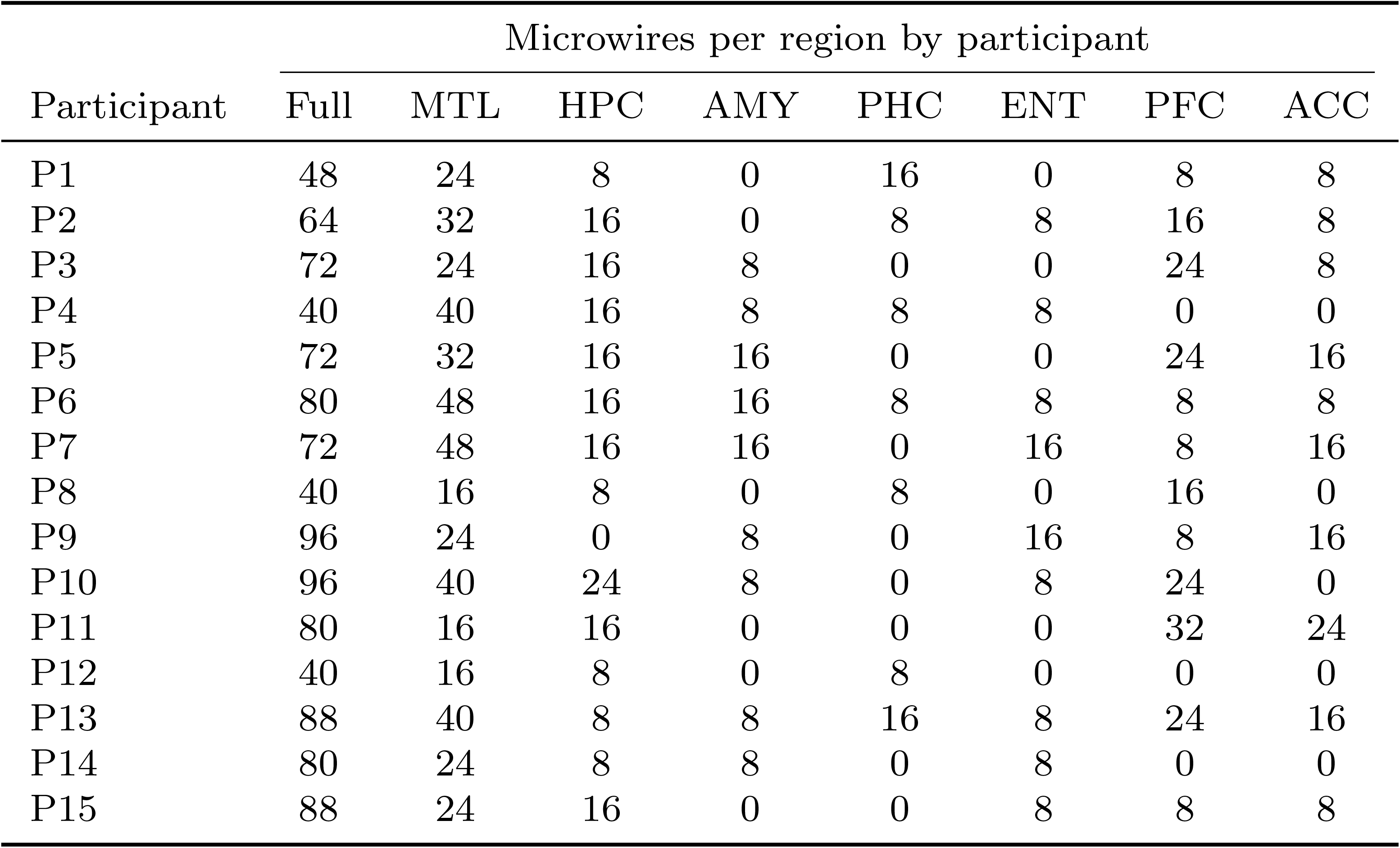
Microwire localizations. Each electrode bundle contains eight wires, which we localize by identifying an artifact on the CT scan visible beyond the tip of the macroelectrode sheath. The CT scan is coregistered with participant T1 and T2 structural MRIs to assess the implanted brain region (see further details in Methods). MTL: medial temporal lobe; HPC: hippocampus; AMY: amygdala; PHC: parahippocampal gyrus; ENT: entorhinal cortex; PFC: prefrontal cortex (includes orbitofrontal, superiorfrontal, and inferiorfrontal); ACC: anterior cingulate cortex. FC: frontal cortex as described in the text represents combined PFC and ACC.

**Table S2.**
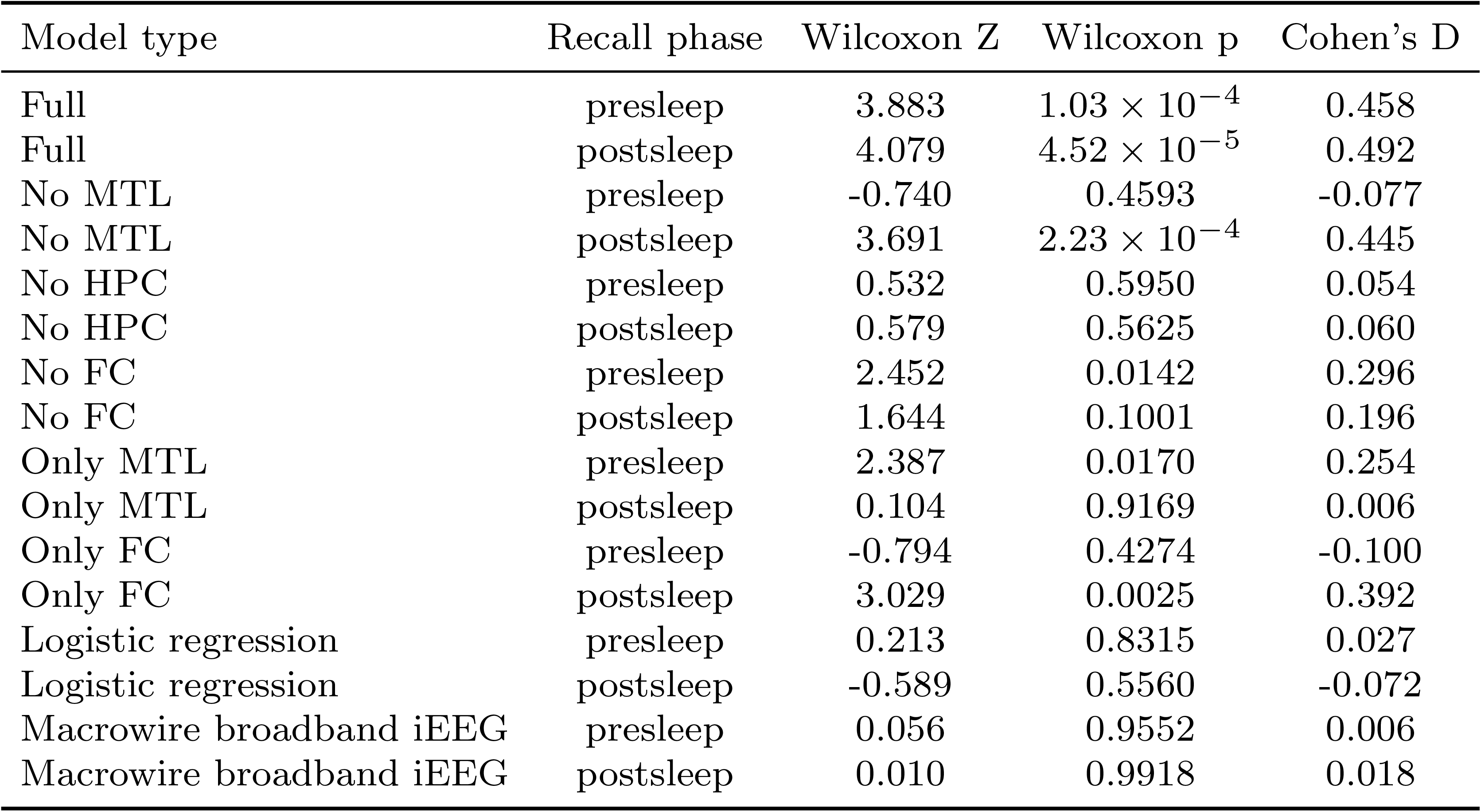
Summary of model statistics. Comparison to chance (50%) for each model using Wilcoxon signed-rank.

**Table S3.**
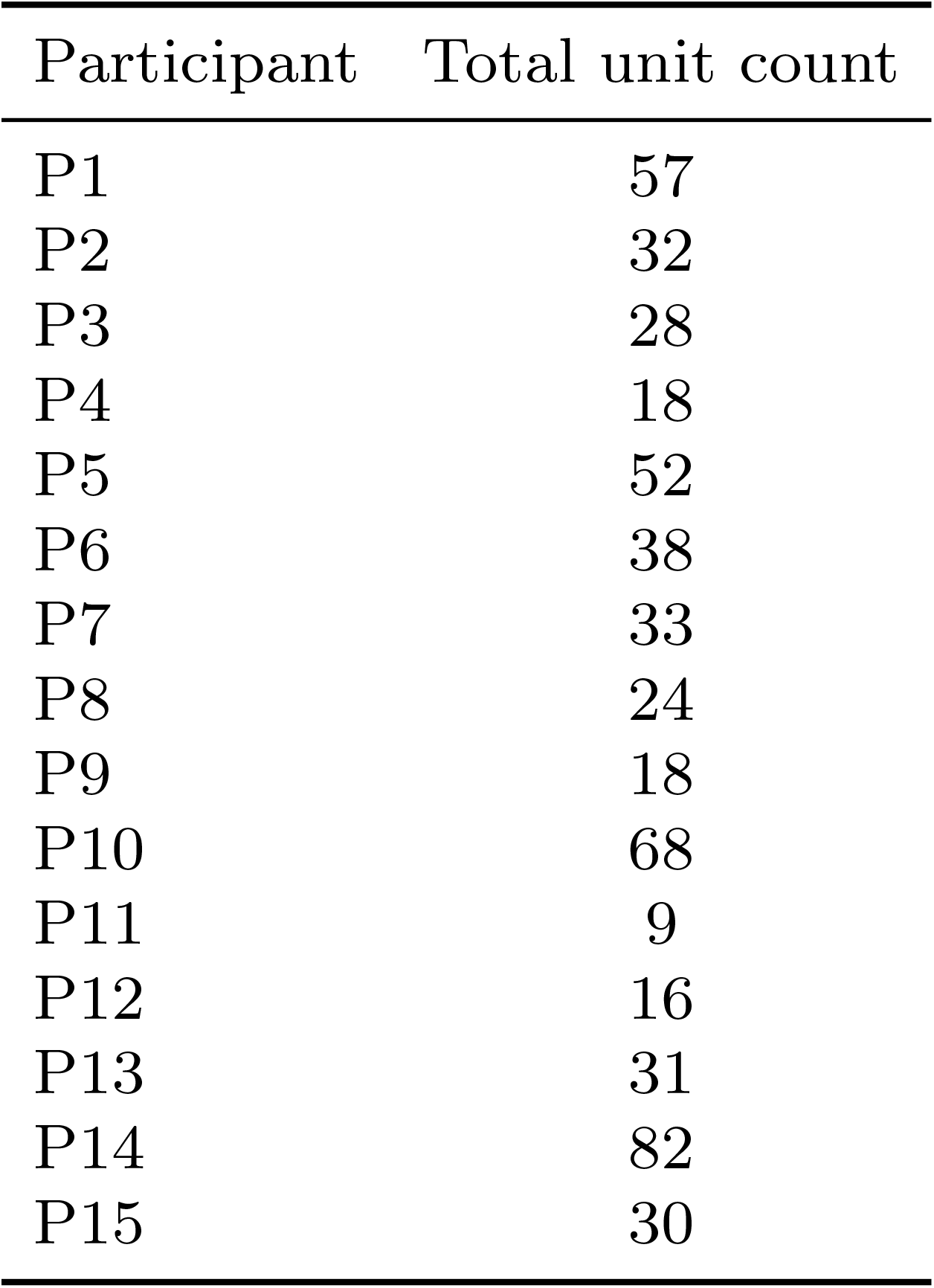
The number of manually curated single- and multi-units with firing rate of > 0.5 spikes/sec over the video watch period for each participant.

**Table S4.** Participant clinical and sleep characteristics.

| ID | Seizure<br>freq. | SOZ | Age | Sex | Medication | Sleep<br>start | Sleep<br>end | NREM<br>(h) | REM/<br>awake<br>(h) |
| --- | --- | --- | --- | --- | --- | --- | --- | --- | --- |
| 1 | 0.3×/mo | Right amygdala | 47 | F | Pregabalin 300 mg qAM and 450 mg qPM, Levetiracetam ER 4000 mg qHS | 22:09 | 07:25 | 5.8 | 1.0 |
| 2 | 1–<br>2×/mo | Bilateral anterior<br>hippocampus | 24 | M | Trileptal 1200 mg BID, Xcopri 200 mg daily | 22:50 | 09:09 | 6.8 | 3.4 |
| 3 | 25×/mo | Left hippocampus | 26 | M | Keppra 3000 mg BID, Fycompa 8 mg QHS, Onfi 40 mg BID, Vimpat 200 mg BID | 23:54 | 09:35 | 5.8 | 0.8 |
| 4 | 1×/mo | Bilateral<br>hippocampus | 25 | M | Keppra 1000 mg BID, Vimpat 200 mg BID, Xcopri 50 mg daily | 02:36 | 08:07 | 4.2 | 0.8 |
| 5 | 1–<br>4×/mo | None | 54 | M | Vimpat 300 mg BID, Lamictal ER 200 mg BID | 00:49 | 08:06 | 2.7 | 1.0 |
| 6 | 2–<br>3×/mo | Right anterior<br>temporal lobe | 41 | F | Xcopri 300 mg daily, Briviact 50 mg BID | 00:21 | 08:04 | 5.2 | 0.9 |
| 7 | 1×/mo | Left anterior MTL | 24 | M | Keppra 2000 mg BID, Aptiom 800 mg QHS, Fycompa 2 mg QHS, Valtoco PRN | 00:52 | 07:52 | 4.9 | 1.4 |
| 8 | 4–<br>6×/mo | Left parietal<br>lesion | 33 | M | Vimpat 200 mg BID, Briviact 150 mg BID, Cenobamate 150 mg daily | 04:28 | 07:46 | 2.3 | 0.3 |
| 9 | 2×/mo | Right MTL | 55 | M | Dilantin 300 mg + 200 mg, Topamax 200 mg BID | 20:09 | 05:59 | 5.4 | 2.3 |
| 10 | 10×/mo | Right MTL, right<br>middle-posterior<br>superior temporal<br>gyrus | 40 | M | Onfi 10 mg daily, Keppra 1500 mg BID, Lamotrigine 200 mg BID | 00:32 | 07:47 | 2.8 | 4.5 |
| 11 | 1–<br>1.5×/mo | Bilateral<br>frontotemporal | 22 | M | Briviact 100 mg BID, Vimpat 150 mg BID | 01:10 | 06:26 | 4.1 | 0.6 |
| 12 | 2×/mo | None | 45 | F | Lamictal ER 250 mg QHS, CBD oil, lion's mane | 22:21 | 08:31 | 6.0 | 2.5 |
| 13 | 3–<br>4×/mo | Right anterior<br>MTL | 21 | F | Briviact 125 mg BID, Lamotrigine 225 mg BID | 01:32 | 08:36 | 3.3 | 2.6 |
| 14 | 2–<br>4×/mo | Left anterior<br>hippocampus and<br>fusiform gyrus | 23 | M | Keppra ER 2250 mg QHS, Zonegran 400 mg QHS, Oxtellar 2400 mg QHS | 23:29 | 08:46 | 5.4 | 2.3 |
| 15 | 5×/mo | Right MTL | 42 | F | Depacote ER 1000 mg + 1500 mg, topiramate 100 mg BID | 20:06 | 05:43 | 4.0 | 1.1 |
| 16 | 1×/mo | Right MTL | 48 | M | Lamictal XR 200 mg BID, Aptiom 800 mg qHS, Briviact 200 mg BID, Xcopri 300 mg daily | N/A | N/A | 0.0 | 0.0 |

